# Acarbose Protects From Central and Peripheral Metabolic Imbalance Induced by Benzene Exposure

**DOI:** 10.1101/2020.03.19.998567

**Authors:** LK. Debarba, A. Mulka, J.B.M. Lima, O. Didyuk, P. Fakhoury, L. Koshko, AA. Awada, K. Zhang, U. Klueh, M. Sadagurski

## Abstract

Benzene is a well-known human carcinogen that is one of the major components of air pollution. Sources of benzene in ambient air include cigarette smoke, e-cigarettes vaping and evaporation of benzene containing petrol processes. While carcinogenic effects of benzene exposure have been well studied, less is known about metabolic effects of benzene exposure. We show that chronic exposure to benzene at low levels induces severe metabolic imbalance in a sex-specific manner, which is associated with hypothalamic inflammation and endoplasmic reticulum (ER) stress. Benzene exposure rapidly activates hypothalamic ER stress and neuroinflammatory responses in male mice, while pharmacological inhibition of ER stress response by inhibiting IRE1α-XBP1 pathway significantly alleviates benzene-induced glial inflammatory responses. Additionally, feeding mice with Acarbose, a clinically available anti-diabetes drug, protected against benzene induced central and peripheral metabolic imbalance. Acarbose imitates the slowing of dietary carbohydrate digestion, suggesting that choosing a diet with a low glycemic index might be a potential strategy for reducing the negative metabolic effect of chronic exposure to benzene for smokers or for people living/working in urban environments with high concentrations of exposure to automobile exhausts.

## 1. Introduction

Benzene, a volatile organic compound (VOC), is known to be one of the major components of air pollution, contributing to both outdoor traffic-related and indoor pollution [1]. Benzene penetrates the body even at low concentrations [2] and was recognized as a priority pollutant by the US Environmental Protection Agency (EPA) [3]. Benzene concentrations vary depending on geographic location [4, 5], duration or route of exposure, and individual susceptibility factors such as age, gender or lifestyle. Various sources of benzene exposure include drinking contaminated water or food, smoking cigarettes, breathing secondhand cigarette smoke, vehicle exhaust or petroleum fuels, vaping e-cigarettes, using paint or detergent products [6-14]. While values of ambient benzene air concentrations in the U.S. have typically been <1 ppm (parts per million), significantly higher levels can be experienced as a result of various environmental factors [15] [16].

For example, benzene exposure levels of ∼50 ppm are particularly relevant to human exposures associated with conventional tobacco cigarettes smoke including passive smoking [17, 18]. While carcinogenic effects of benzene exposure have been well studied, less is known about metabolic effects of benzene exposure with particular relevance to chronic diaseses such as Type 2 diabetes mellitus (T2DM) [19].

T2DM has become a global pandemic resulting in a reduced quality of life due to comorbidity and premature mortality [20-22]. A recent study demonstrated an association between levels of urinary benzene metabolites in elderly adults living in polluted areas and insulin resistance (IR) [23]. Additionally, benzene exposure and presence of urinary benzene metabolites are associated with increased cardiovascular risk and dyslipidemia in both smokers and nonsmokers, suggesting that benzene may be linked to cardiovascular injury, even in the absence of exposure to other tobacco smoke constituents [16]. Benzene exposure during pregnancy is strongly associated with gestational diabetes among women belonging to underrepresented minorities in the US [24]. T2DM was found to significantly increase in the self-reported Benzene Sub-registry in the NHIS survey [25]. An association between urinary benzene metabolites and risk of IR in children and adolescents provided an indication that benzene exposures, even at low levels, might be associated with IR [26]. Young mice exposed to volatile benzene develop IR and inhibition of insulin signaling in the liver and skeletal muscle, associated with activation of inflammatory signaling pathway [27]. Together, these data imply that benzene exposure may be a significant and heretofore overlooked risk factor for IR and subsequent T2DM development. Herein we demonstrate that exposure to 50 ppm of benzene leads to severe hyperglycemia and hyperinslulinemia without any adverse toxicological effects observed.

The hypothalamus is critical in sensing circulating metabolic signals and regulating neural pathways to control peripheral glucose metabolism [28-30]. Under pathological conditions such as obesity and diabetes, hypothalamic neurons are interrupted due to the induction of metabolic inflammation, mediated by endoplasmic reticulum (ER) stress response and proinflammatory pathways [29, 31]. In rodent models of obesity, an accumulation of reactive, pro-inflammatory microglia and astrocytes in the hypothalamus are evident within 24 hours of high-fat diet (HFD) onset, prior to substantial weight gain [32]. Inhalation of PM with diameters of 2.5 micrometers (PM_2.5_) results in hypothalamic inflammation and central inhibition of inflammatory NF-kB signaling prevents PM_2.5_ mediated peripheral inflammation and exacerbation of T2DM [33]. These findings suggest that hypothalamic function may be of fundamental importance in the regulation of inflammatory changes that may contribute to the development of air pollution induced T2DM. In the present study, we demonstrate the role of hypothalamic mechanisms as targets for benzene diabetogenic effects on metabolism.

Benzene is present in different concentrations in all human day to day activities. Since exposure to benzene is unavoidable, interventions that can potentially mitigate the adverse health effects of benzene exposure are needed. Acarbose (ACA) is an inhibitor of intestinal α-glucosidase which slows carbohydrate digestion [34]. ACA has been used clinically to prevent postprandial hyperglycemia for many years, and has been used to treat patients with T2DM or glucose intolerance [35]. Similarly to metformin, ACA has beneficial effects on body weight, onset of diabetes and cardiovascular risk [36, 37]. In mice, ACA effect replicates some aspects of diet or caloric restriction associated with increased lifespan [38-41], and with inhibition of age-associated hypothalamic inflammation in a sex-specific manner [42]. Our findings indicate that hypothalamic ER stress and hypothalamic inflammation might be critical for benzene diabetogenic effects, while ACA feeding, protects against benzene induced hypothalamic inflammation, ER stress and peripheral metabolic imbalance.

## 2. Materials and Methods

### 2.1. Benzene exposure

7 weeks-old male and female C57BL/6 were purchased from The Jackson Laboratory. The mice in inhalation chambers using FlexStream™ automated Perm Tube System (KIN-TEK Analytical, Inc) were exposed to benzene concentration of 50 ppm for 6h/day for 4 weeks (Supplementary Figure 1A). FlexStream™ automated Perm Tube System allows creating precision gas mixtures. This unit provides a temperature-controlled permeation tube oven, dilution flow controls, and front panel touch-screen interface. Mixtures are produced by diluting the miniscule flow emitted from Trace Source™ permeation tubes with a much larger flow of inert matrix gas, typically nitrogen or zero air. Control animals were breathing regular filtered room air. All mice were provided with water *ad libitum* and housed in temperature-controlled rooms on a 12-hour/12-hour light-dark cycle.

### 2.2. Diets

All mice were provided *ad libitum* access to standard chow diet (Purina Lab Diet 5001), high-fat diet (HFD) (Research Diets, #D12451) or Acarbose diet, as before [38, 39, 42]. Acarbose diet (Spectrum Chemical Mfg. Corp., Gardena, CA, product # A3965, CAS # 56180-94-0) [38] was kindly provided by Randy Strong laboratory (UT San Antonio, Texas) as part of NIA’s Intervention Testing Program (ITP). Acarbose was fed continuously at a concentration of 1,000 mg of Acarbose per kilogram of diet (ppm) to mice while exposed to benzene in the chambers, as indicated.

### 2.3. Metabolic Analysis

Lean and fat body mass were assessed by a Bruker minispec LF 90II NMR-based device. Blood glucose levels were measured on random-fed or overnight-fasted animals in mouse-tail blood using Glucometer Elite (Bayer). Intraperitoneal glucose tolerance tests were performed on mice fasted for 6 hours. Animals were then injected intraperitoneally with D-glucose (2 g/kg) and blood glucose levels were measured as before [71]. For an insulin tolerance test, animals fasted for 5 hours received an intraperitoneal injection of human insulin (0.5 units/kg; Novo Nordisk). Blood insulin was determined on serum from tail vein bleeds using a Rat Insulin ELISA kit (Crystal Chem. Inc.). Blood levels of low-density lipoprotein cholesterol (LDL-C), Non-Esterified Fatty Acid (NEFA) and triglycerides were determined by ELISA kits from Fujifilm Wako Diagnosis.

### 2.4. Perfusion and Histology

Mice were anesthetized (IP) with avertin and transcardially perfused with phosphate-buffered saline (PBS) (pH 7.5) followed by 4% paraformaldehyde (PFA). Brains were post-fixed, sank in 30% sucrose, frozen in OCT medium and then sectioned coronally (30 µm) using a Leica 3050S cryostat. Four series were collected and stored at –80°C in cryo protectant until processed for immunohistochemistry as previously described [72]. For immunohistochemistry, free-floating brain sections were washed in PBS, blocked using 3% normal donkey serum (NDS) and .3% Triton X-100 in PBS and then stained with a primary antibody overnight in blocking buffer. For IRE-1α, immunostaining, sections were pretreated for 20 min in 0.5% NaOH and 0.5% H2O2 in PBS, followed by immersion in 0.3% glycine for 10 min and placed in 0.03% SDS for 10 min then stained with primary IRE1-1α (anti-rabbit, Cell Signaling Tech, 1:100) antibody overnight. Other floating brain sections were washed with PBS several times; blocked for 1h in 0.3% Triton X-100 with 3% normal donkey serum in PBS; and then stained with the following primary antibodies overnight: GFAP (anti-rabbit, Millipore, 1:1000), Iba1 (anti-goat, Abcam, 1:1000). All floating brain sections were washed with PBS several times; and incubated with the anti-rabbit, anti-gout Alexa Fluor 488 and/or 568 (Invitrogen, 1:200) secondary antibody for 2h. Sections were mounted onto Superfrost Plus slides (Fisher Scientific, Hudson, NH) and coverslips added with ProLong Antifade mounting medium (Invitrogen, Carlsbad, CA). All images were visualized with Nikon 800 fluorescent microscope using Nikon imaging DS-R12 color cooled SCMOS, version 5.00.

### 2.5. Morphology assessment

Immunofluorescence images were taken using a multiphoton laser-scanning microscope (LSM 800, ZEISS) equipped with a 63X objective for the arcuate nucleus of the hypothalamus. Stacks of consecutive images taken at 0.7-μm intervals were sequentially acquired, and 27 optical sectioning, along the optical axis (z-axis), were reconstructed to 3D using Fiji-Image J. Measurements of the area of soma size were analyzed using Fiji-Image J software. Measurements of process extensions were analyzed using the Sample Neurite Tracer of Fiji-Image J software [73, 74].

### 2.6. Glial culture

The whole-brain from 1-2 day-old neonatal mice was collected, without the meninges, placed in HBSS modified medium without calcium or magnesium (Gibco 14190144) and processed according to the manufacturer’s instructions from Miltenyi Biotech. The supernatant was discarded, and the pellet was re-suspended in 1–2 mL of Dulbecco’s Modified Eagle Medium/Nutrient Mixture F-12, no phenol red (DMEM/F-12), (Gibco 21041025) enriched with 10% inactivated fetal bovine serum (FBS) (Thermo Fischer 10082147) and 1% Antibiotic-Antimycotic (Thermo Fischer 15240062). After homogenization, the cells were grown in Nunc™ Cell Culture Treated Flasks with Filter Caps, (Thermo Fischer, 178905) containing 10 mL of the medium [DMEM/F-12; 10% FBS; 1% Antibiotic-Antimycotic] in a 5% CO_2_ incubator (Galaxy 170R, Eppendorf) at 37°C. The cells were maintained in culture for a period of 8–10 days, with a partial replacement of the incubation medium (70%) every 48–72h. At 80% confluence, the cells were incubated in a shaker at 200 rpm for 2 h at 37°C to separate the oligodendrocytes and neurons from the glial cell culture. The cells in suspension were discarded, and the adhered cells were washed with 1 mL of Dulbecco’s Phosphate-Buffered Saline, without calcium or magnesium (DPBS), (Gibco 14190144). Then, 5 mL of 1x trypsin (Thermo Fischer 15400054) were added, and the cells were maintained for 15 min in the incubator (5% CO_2_ at 37°C). After the exposure to benzene or an inhibitor of IRE1α, STF-083010 (from Zhang lab, WSU) the cells were washed with 1 ml of cold DPBS and immediately harvested with 1 ml of trizol, following for RNA extraction as described below. STF-083010 is an inhibitor of the IRE1α -XBP1 pathway, blocking IRE1 endonuclease activity without affecting its kinase activity after ER stress both *in vitro* and *in vivo [75, 76]*.

### 2.7. RNA extraction and qPCR

Hypothalamus was carefully dissected using Brain Matrices (Braintree Scientific, Braintree, MA). Total RNA was extracted from tissues with Trizol (Gibco BRL) and 1000 ng of RNA was used for cDNA synthesis using iscript cDNA kit (Bio-Rad Laboratories Inc.). Quantitative real-time PCR was performed using the Applied Biosystems 7500 Real-Time PCR System. The following SYBER Green Gene Expression Assays (Applied Biosystems) were used in this study Table 1 Each PCR reaction was performed in triplicate. Lack of reverse transcription (-RT) was used as a negative control for contaminating DNA detection. Water instead of cDNA was used as a negative control, and the housekeeping gene ß-actin was measured in each cDNA sample. Gene transcripts in each sample were determined by the ΔΔCT method. For each sample, the threshold cycle (CT) was measured and normalized to the average of the housekeeping gene (ΔCT). The fold change of mRNA in the unknown sample relative to the control group was determined by 2-ΔΔCT. Data are shown as mRNA expression levels relative to the control group.

### 2.8. Statistical analysis

Results are expressed as the mean ± standard error and were analyzed using Statistica software (version 10). An analysis of t-test was used when compared only benzene treatment versus control, variance (ANOVA) with repeated measurements was used to analyze body weight, GTT, and ITT. Other parameters were analyzed by two-way ANOVA (treatment and gender), (Treatment and Inhibitor) or (treatment and diet) All data analyzed with a Two way ANOVA were further analyzed with Newman-Keuls post hoc analysis The level of significance (α) was set at 5%.

### 3. Results

### 3.1. Benzene exposure induces hyperglycemia and hyperinsulinemia in a sex-specific manner

The effect of 50 ppm benzene exposure in mice was assessed by direct inhalation for 6 hr/day, 5 days/wk for a period of 4 weeks. Supplementary Figure 1A depicts a schematic of the exposure protocol in benzene inhalation chambers. This adapted regimen for benzene inhalation resembles human benzene exposure levels in highly polluted gas stations, or exposure to tobacco smoke or e-cigarette vapors aerosols [12, 43, 44]. Under these conditions, benzene exposure had no effect on body weight or body composition (fat and lean mass) in chow-fed male or female mice (Figure 1 A and B, and Supplementary Figure 1B and C). Moreover, there were no significant changes in white blood cell counts in benzene exposed animals versus control animals breathing room air (Supplementary Figure 1D), while levels of the urine benzene metabolite, t,t-MA were significantly increased in benzene exposed mice (Supplementary Figure 1E). Additionally, the gene expression of P450 2E1 (*Cyp2e1*), a metabolizing enzyme involved in responses to benzene [45] was significantly increased in the liver of benzene exposed mice (Supplementary Figure 1G). Despite normal fasting glucose levels, fasting insulin concentrations were significantly elevated and glucose tolerance of benzene exposed male mice was significantly impaired (Figure 1C and D). In contrast, female mice were completely resistant to the negative metabolic consequences of chronic benzene exposure (Figure 1B, E and F). Insulin tolerance was not impaired in both benzene-exposed male and female mice (Supplementary Figure 2A and B). When kept on a high-fat diet (HFD) for 8 weeks while exposed to benzene, both male and female mice gained the same amount of weight, lean and fat mass, and demonstrated glucose intolerance similar to HFD-fed control littermates (Figure 1 G, H and Supplementary Figure 3). Thus, an obesogenic diet initiated in parallel with benzene exposure does not exacerbate benzene effects on glucose metabolism. We also detected an increase in expression of genes associated with gluconeogenesis (*G6pc and Ppck1*) and lipid and fatty acids synthesis (*Srebp1, Srepb2, Abca1, Cpt1, and Acc*) in livers from benzene-exposed male mice as compared to control mice, but not in female mice (Figure 2A and B). Additionally, serum levels of triglyceride, cholesterol and free fatty acids were significantly elevated in benzene-exposed male mice (Figure 2C-E). Benzene-exposed mice exhibited an increase in hepatic inflammatory genes *Il1* and *Il6* (Figure 2F and G) in both sexes, suggesting that benzene induced metabolic imbalance in male mice does not depend on peripheral hepatic inflammation. Together these findings indicate that benzene exposure alters glucose homeostasis and influences peripheral lipid metabolism in a sex-specific manner independent of hepatic inflammatory responses.

**Figure 1.**
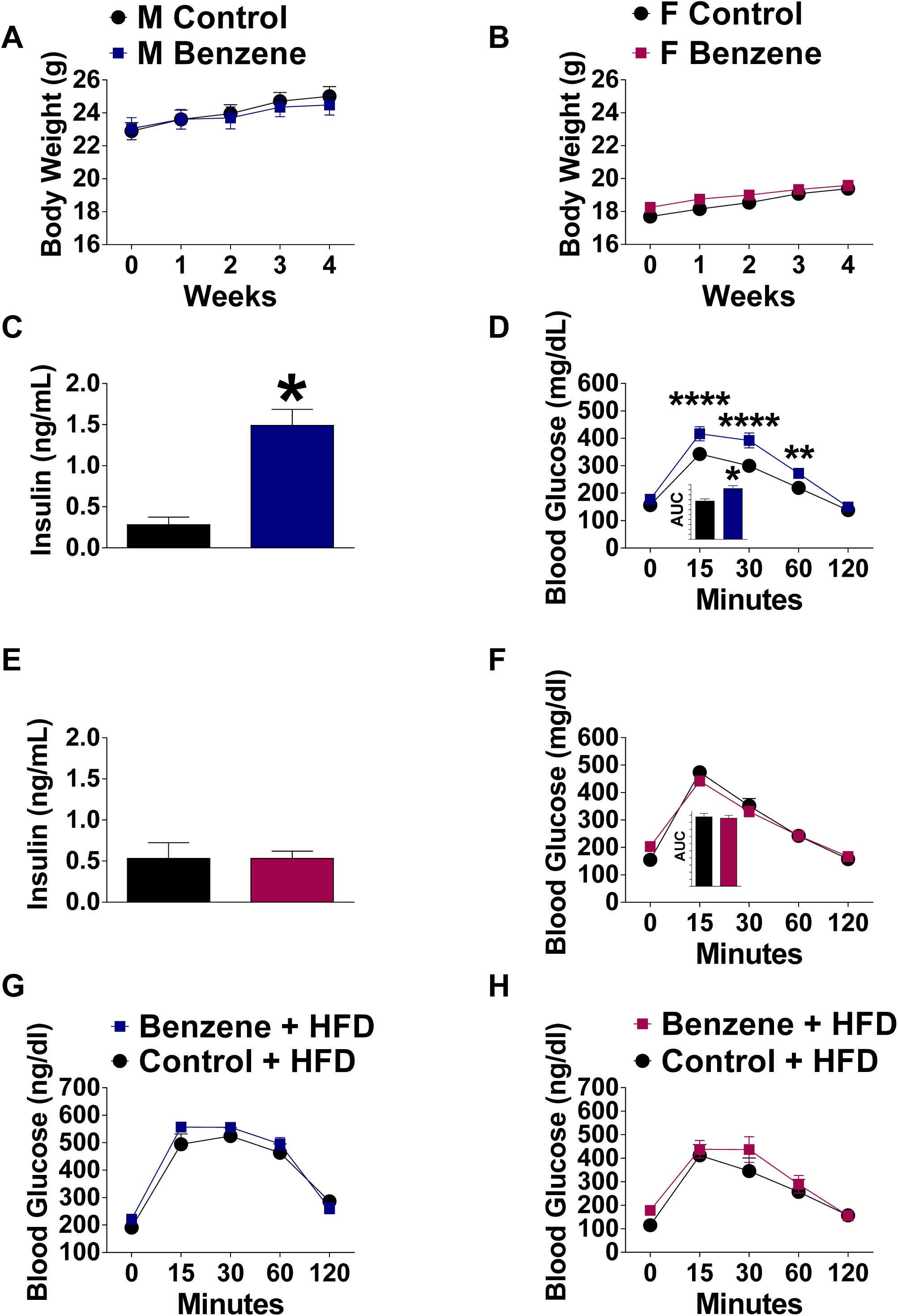
Chronic benzene exposure promotes hyperglycemia and hyperinsulinemia in male but not in female mice. Body weight in (**A**) males (blue) and (**B**) females (magenta); (**C**) fasted insulin serum levels (*t* = 15.82, *p* < 0.0001), (n=8) and (**D**) Glucose tolerance test (GTT) (effect of treatment: *F*_1.9_ = 24.83, *p*< 0.0007, *η*^2^ = 73.39%; effect of time: *F*_4.36_ = 229.25, *p*< 0.0001, *η*^2^ = 96.22%; and treatment x time interaction *F*_4.36_ = 16.66, *p* < 0.0001, *η*^2^ = 65.05; and area under curve (AUC) (*t* = 7.269, *p* < 0.0001), %) (n = 15-16 mice per group). **(E)** Fasted insulin serum levels (**F**) GTT and AUC of female mice (**G**) GTT in HFD-fed male and (**H**) HFD-fed female mice exposed to 50 ppm benzene. Data is expressed as the mean ± SEM. Repeated measures ANOVA were further analyzed with Newman-Keuls post hoc analysis (* = *vs* control; \**p* < 0.05; \*\**p* < 0.01; \*\*\**p* < 0.001; \*\*\*\**p* < 0.0001).

**Figure 2.**
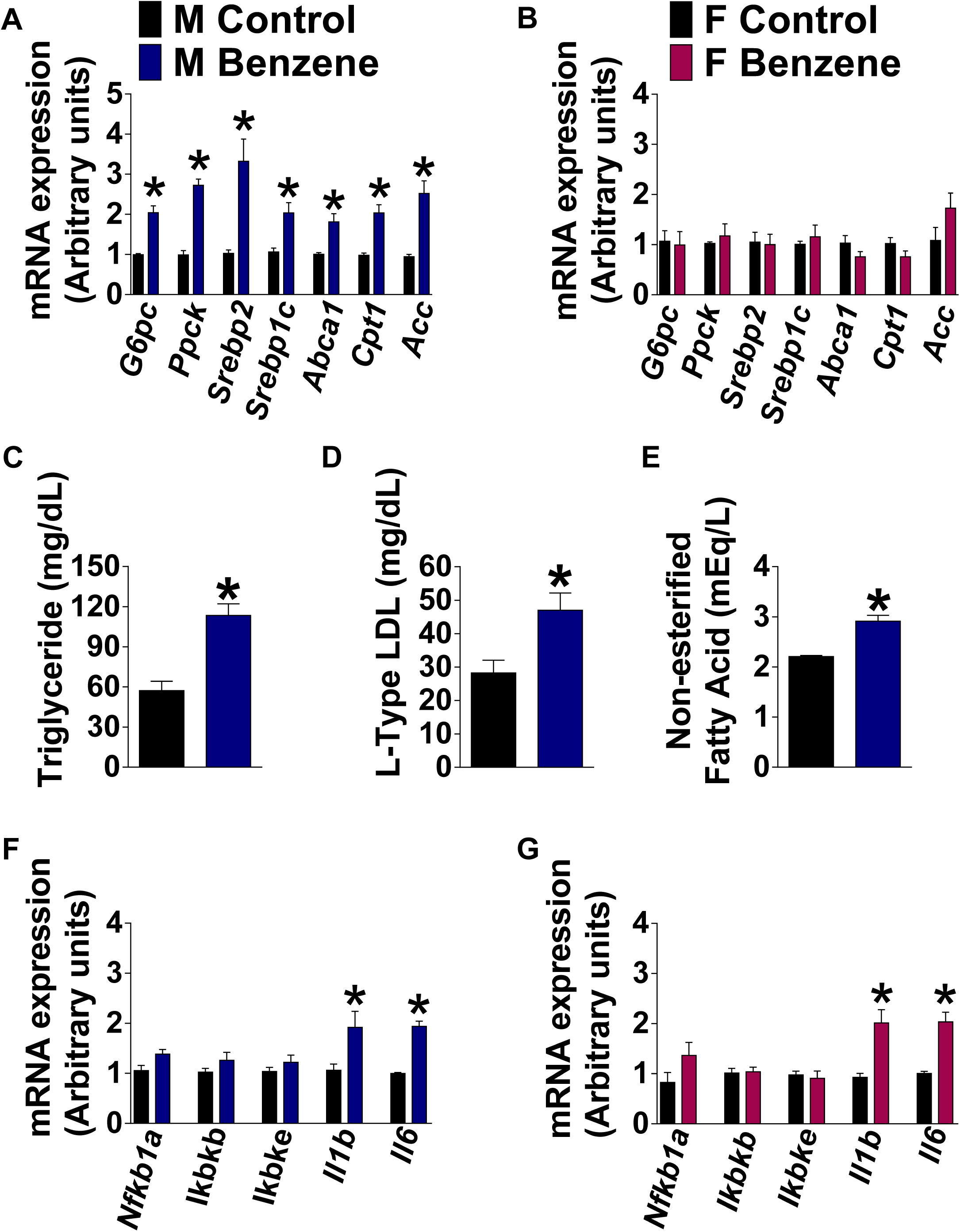
Effect of chronic benzene exposure on hepatic gene expression. (**A**) qPCR of hepatic genes associated with gluconeogenesis *G6pc* (*t* = 5.948, *p* < 0.0003) and *Ppck* (*t* = 9.719, *p* < 0.0001); lipid and fatty acids synthesis, *Srepb2* (*t* = 4.090, *p* < 0.0009), *Srebp1c* (*t* = 3.497, *p* < 0.0036), *Abca1* (*t* = 3.889, *p* < 0.0013), *Cpt1* (t=5.003, *p* < 0.0001), and *Acc* (*t* = 4.995, *p* < 0.0001), of males and (**B**) females, (n=8-9 per each group); Blood serum levels of (**C**) triglyceride (mg/dL) (*t* = 5.037, *p* < 0.0005); (**D**) Low-density lipoprotein (LDL) (mg/dL) (*t* = 2.915, *p* < 0.0015) and (**E**) Non-esterified fatty acid (FFA) (mEq/L) (*t* = 5.834 df=10, *p* < 0.0002) of males, (n=6). qPCR of hepatic inflammatory genes (*Nfkb1a, Ikbkb, Ikbke, Il1b, Il6*) of (**F**) males and (**G**) females, (n=7 for each group). *For Il1b* (males: *t=8.439, p* < 0.0001; females: *t=3.984, p* < 0.0018) and *Il6* (males: *t* = 8.773, *p* < 0.0001; females: *t* = 5.158, *p* < 0.0002). Data is expressed as the mean ± SEM. (* = *vs* control; \**p* < 0.05)

### 3.2. Benzene induced hypothalamic inflammation

To assess the effect of benzene exposure on hypothalamic inflammation, we examined the numbers of microglia and astrocytes in male and female mice exposed to benzene for 4 weeks. Astrogliosis is correlated with increased expression of the glial fibrillary acidic protein (GFAP) [32]. Accordingly, 4-weeks of benzene exposure resulted in a significant increase in GFAP^+^ astrocytes in the different areas of the medial basal hypothalamus (MBH) such as the arcuate nucleus (ARC), the ventromedial (VMH), the dorsomedial (DM) hypothalamus as compared to control male mice (Figure 3A and B). Consistent with previous studies [42, 46, 47], control females demonstrated a significantly lower number of immunostained GFAP^+^ astrocytes in the MBH compared to control male mice (Figure 3A and B). Additionally, using immunostaining for the microglia-specific ionized calcium-binding adaptor molecule 1 (Iba1), we observed that the number of microglial cells in the MBH is significantly higher in benzene exposed male as compared to control mice (Figure 3C and D). Strikingly, females exposed to benzene demonstrated similar numbers of GFAP^+^ astrocytes and Iba^+^ microglia cells to controls (Figure 3A-D), suggesting the protective gender effect in exposure to benzene. Similarly, the expression levels of inflammatory genes *Ikbkb, Ikbke, Tnf, Il1b*, and *Il6* measured from the microdissected ARC were significantly elevated in benzene exposed male mice, indicating an inflammatory status within the ARC, and the expression of *Cyp2e1* was elevated (Figure 3E), supporting the benzene-induced activation of CYP2E1 in glial cells[48].

**Figure 3.**
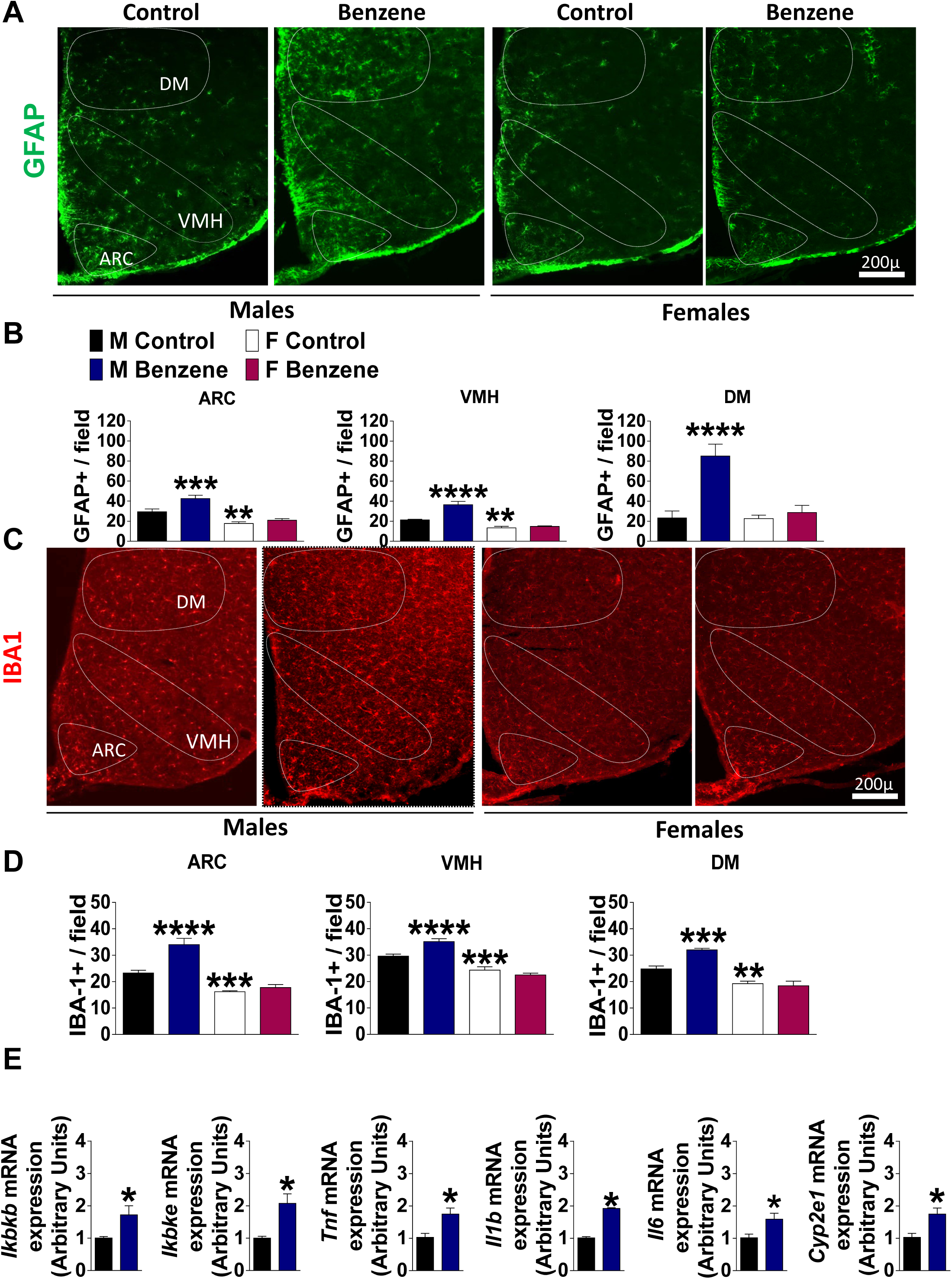
Chronic benzene exposure-induced hypothalamic gliosis. (**A**) Representative images of astrocytes identified by immunofluorescent detection of GFAP protein (**B**) number of astrocytes in the ARC (effect of treatment: *F*_1.22_ = 11.54, *p*< 0.0025, *η*^2^ = 13.30%; effect of gender: *F*_1.22_ = 48.83, *p*< 0.0001, *η*^2^ = 56.27%; and treatment x gender interaction *F*_1.22_ = 4.40, *p* < 0.0001, *η*^2^ = 5.07%), VMH (effect of treatment: *F*_1.23_ = 21.92, *p*< 0.0001, *η*^2^ = 16.91%; effect of gender: *F*_1.23_ = 69.45, *p*< 0.0001, *η*^2^ = 53.59%; and treatment x gender interaction *F*_1.23_ = 15.21, *p* < 0.0007, *η*^2^ = 11.74%) and DMH (effect of treatment: *F*_1.23_ = 20.39, *p*< 0.0001, *η*^2^ = 28.46%; effect of gender: *F*_1.23_ = 14.46, *p*< 0.0009, *η*^2^ = 20.19%; and treatment x gender interaction *F*_1.23_ = 13.78, *p* < 0.0011, *η*_2_ = 19.24%), (n=6-8); (**C**) Representative images of microglia identified by immunofluorescent detection of Iba-1 protein (**D**) quantification of microglia in the ARC (effect of treatment: *F*_1.24_ = 22.35, *p*< 0.0001, *η*^2^ = 16.02%; effect of gender: *F*_1.24_ = 80.79, *p*< 0.0001, *η*^2^ = 57.90%; and treatment x gender interaction *F*_1.24_ = 12.38, *p* < 0.0018, *η*^2^ = 8.87%), VMH (effect of treatment: *F*_1.23_ = 40.92, *p*< 0.0248, *η*^2^ = 4.39%; effect of gender: *F*_1.23_ = 85.31, *p*< 0.0001, *η*^2^ = 64.98%; and treatment x gender interaction *F*_1.23_ = 17.20, *p* < 0.0003, *η*^2^ = 13.10%), and DMH (effect of treatment: *F*_1.23_ = 7.82, *p*< 0.0102, *η*^2^ = 6.94%; effect of gender: *F*_1.23_ = 69.68, *p*< 0.0001, *η*^2^ = 61.83%; and treatment x gender interaction *F*_1.23_ = 12.20, *p* < 0.0019, *η*^2^ = 10.82%), (n=6-8 per group) (**E**) qPCR of inflammatory genes: *Ikbkb* (*t=2.503, p* < 0.0464), *Ikbke* (*t=3.577, p* < 0.0117), *Tnf* (*t= 3.214, p* < 0.0183), *Il1b* (*t* = 18.70, *p* < 0.0001), *Il6* (*t* = 2.700, *p* < 0.0356) and *Cyp2e1* (*t*=3.214, *p* < 0.0183), (n=5 per group), analyzed from the ARC of males. Data is expressed as the mean ± SEM and analyzed with a Two-way ANOVA followed with Newman-Keuls post hoc analysis, (* = vs male control; \**p* < 0.05; \*\**p* < 0.01; \*\*\**p* < 0.001; \*\*\*\**p* < 0.0001).

We next explored the effect of acute 6 hours benzene exposure on hypothalamic inflammation. Figure 4 demonstrates that acute benzene exposure triggered significant reactive gliosis in the MBH (Figure 4 A-D) and an increase in inflammatory genes in the ARC of benzene-exposed male mice as compared to control animals (Supplementary Figure 4). Additionally, acute benzene exposure induced rapid changes in the astroglial morphology. Astrocytic soma area and the total volume of astrocytic processes were drastically increased in the ARC of benzene exposed male mice, as analyzed by confocal microscopy followed by 3D morphology reconstruction analysis (Figure 4 E-G). Benzene induced hypothalamic inflammatory responses were not observed in female mice (Figure 4 A-D). These data suggest that benzene induced hypothalamic inflammation may precede and contribute to metabolic imbalance observed in benzene exposed animals, and further provide evidence for the male vulnerability to severe neuroinflammation upon benzene exposure.

**Figure 4.**
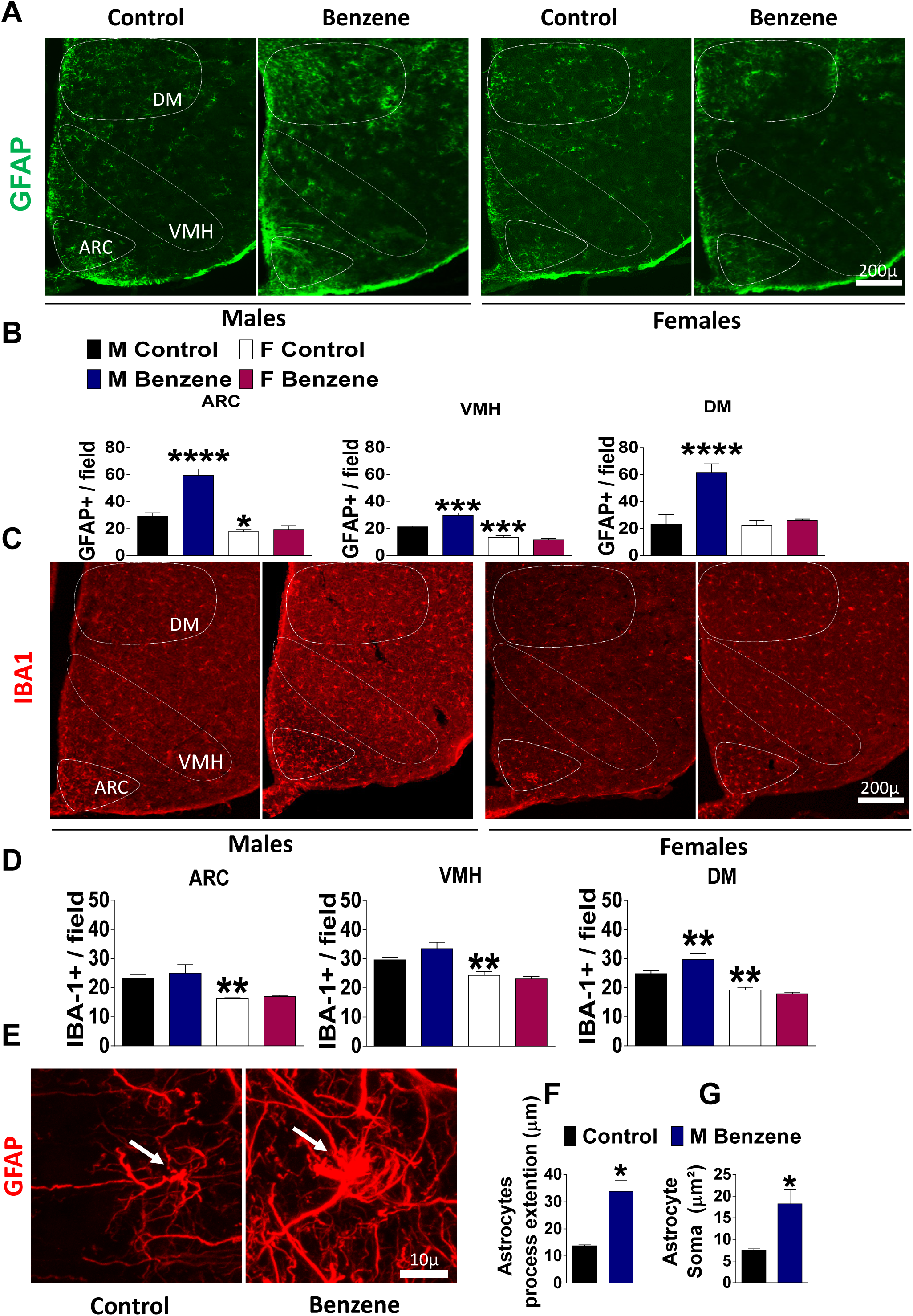
Acute benzene exposure-induced hypothalamic gliosis. (**A**) Representative images of astrocytes identified by immunofluorescent detection of GFAP protein (**B**) Quantification of astrocytes in the ARC (effect of treatment: *F*_1.22_ = 29.02, *p*< 0.0001, *η*^2^ = 19.26%; gender: *F*_1.22_ = 76.48, *p*< 0.0001, *η*^2^ = 50.77%; and treatment x gender interaction *F*_1.22_ = 23.15, *p* < 0.0001, *η*^2^ = 15.37%), VMH (treatment: *F*_1.23_ = 6.06, *p*< 0.0217, *η*^2^ = 4.34%; gender: *F*_1.23_ = 95.66, *p*< 0.0001, *η*^2^ = 68.58%; and treatment x gender interaction *F*_1.23_= 14.77, *p* < 0.0001, *η*^2^ = 10.59%), and DMH (treatment: *F*_1.23_ = 16.62, *p*< 0.0004, *η*^2^ = 26.05%; gender: *F*_1.23_ = 12.55, *p*< 0.0017, *η*^2^ = 19.67%; and treatment x gender interaction *F*_1.23_= 11.63, *p* < 0.0023, *η*^2^ = 18.23%); (n=6-9 per each group) **(C)** Representative images of microglia identified by immunofluorescent detection of Iba-1 protein **(D)** Quantification of microglia in the ARC (effect of gender: *F*_1.24_ = 30.10, *p*< 0.0001, *η*^2^ = 54.66%), VMH (effect of gender: *F*_1.23_ = 32.09, *p*< 0.0001, *η*^2^ = 54.05%) and DMH (effect of gender: *F*_1.23_ = 50.06, *p*< 0.0001, *η*^2^ = 61.33%; and treatment x gender interaction *F*_1.23_= 6.44, *p* < 0.0184, *η*^2^ = 7.89%) (n=6-9); (**E**) Representative images of astrocyte morphology in the ARC of male mice (**F**) astrocyte process extension (*t=5.123, p* < 0.0009), and (**G**) area of soma (*t* = t=3.154, *p* < 0.0135), (n=6). Data was analyzed with a Two-way ANOVA followed by Newman-Keuls post hoc analysis, (* = vs male control; *p < 0.05; **p < 0.01; ***p < 0.001; ****p < 0.0001).

### 3.3. Benzene induced hypothalamic ER stress

The endoplasmic reticulum (ER) stress and unfolded protein response (UPR) are related to the induction of proinflammatory cytokines [49]. Chronic benzene-exposure significantly increased the immunoreactivity of ER stress-induced inositol-requiring enzyme IRE1α, the primary UPR transducer [50], in the ARC of male mice compared to controls (Figure 5A and B). Importantly, acute exposure to benzene for 6 hours was sufficient to induce a robust increase in the immunoreactivity of IRE1α (Figure 5C and D) and gene expression of *Ire1α*, X-box binding protein 1(*Xbp1*), and transcription factor C/EBP homologous protein (*Chop*) in the ARC of male mice (Figure 5E). To further study the role of the IRE1α-UPR pathway in benzene-induced acute gliosis, a specific IRE1α RNase inhibitor STF-083010 was used to suppress IRE1α signaling in primary glial cells exposed to benzene [50 nM]. Under ER stress, IRE1α RNase processes Xbp1 mRNA, and spliced Xbp1 mRNA (Xbp1s) encodes a potent transcription factor that functions as a major mediator of the IRE1-UPR pathway. It has been established that Xbp1s can potentially drive pro-inflammatory response by promoting expression of the pro-inflammatory cytokines TNF-α, IL6 and IL1β in macrophages [51] [52]. As shown in Figure 5F, exposure of primary glial cells to benzene induced the upregulation of *Xbp1s*, and inflammatory *IL-1β* and *TNF-α* gene expression in a manner similar to LPS[53]. Treatment with STF-083010 resulted in significant inhibition of benzene-induced *Xbp1s* expression, followed by reduced expression of *TNF-α* and *IL-1β* (Figure 5F), indicating that benzene-induced ER stress might induce inflammatory responses in glial cells mediated through the IRE1α-XBP1 UPR branch.

**Figure 5.**
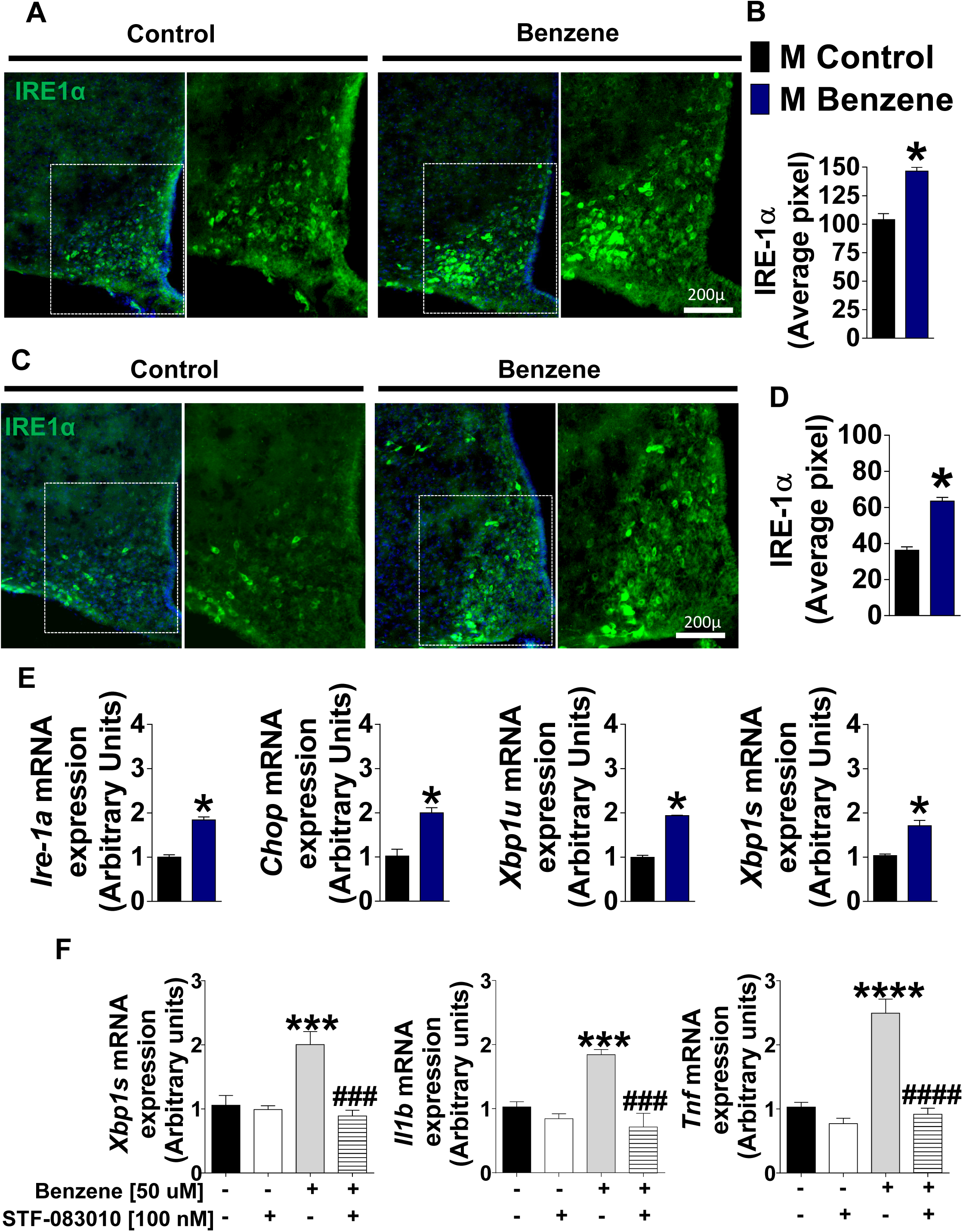
Benzene exposure-induced hypothalamic ER stress. **(A**) Representative images of IRE1α immunoreactivity in the ARC of chronic benzene exposed male mice (**B**) Quantification of IRE1α immunoreactivity (*t* = 6.663, *p* < < 0.0001, *n* = 8). (**C**) Representative images of IRE1α in the ARC of acute (6 hours) benzene exposed male mice (**D**) Quantification of IRE1α immunoreactivity (*t* = 9.683, *p* < < 0.0001, *n* = 8). (**E**) Gene expression of ER stress markers *Ire-1a* (*t* = 9.477, *p* < < 0.0001), *Chop* (*t* = 5.193, *p* < 0.0035), *Xbp1*_U_ (*t* = 20.13, *p* < 0.0001) and spliced, *Xbp*1_S_ (*t* = 4.459, *p* < 0.0066), (n=4) (**F**) Primary glial cell culture treated with benzene (50 nM) and IRE1α inhibitor, STF-083010 (effect of treatment: *F*_1.17_ = 9.20, *p*< 0.0075, *η*^2^ = 15.75%; effect of inhibitor: *F*_1.17_ = 18.08, *p*< 0.0005, *η*^2^ = 30.94%; and treatment x inhibitor interaction *F*_1.17_= 14.15, *p* < 0.0015, *η*^2^ = 24.22%), *Il1b* (effect of treatment: *F*_1.17_ = 9.20, *p*< 0.0075, *η*^2^ = 11.31%; effect of inhibitor: *F*_1.17_ = 28.27, *p*< 0.0001, *η*^2^ = 45.85%; and treatment x inhibitor interaction *F*_1.17_ = 14.65, *p* < 0.0013, *η*^2^ = 21.67%), and *Tnfα* (effect of treatment: *F*_1.17_ = 39.36, *p*< 0.0001, *η*^2^ = 29.37%; effect of inhibitor: *F*_1.17_ = 51.07, *p* < 0.0001, *η*^2^ = 38.16%; and treatment x inhibitor interaction *F*_1.17_ = 26.42, *p* < 0.0001, *η*^2^ = 19.75%), (*n* = 6-5). Data is expressed as the mean ± SEM and analyzed with a Two-way ANOVA followed by Newman-Keuls post hoc analysis, (* = *vs* control; *p < 0.05; **p < 0.01; ***p < 0.001; ****p < 0.0001), (# = *vs* benzene, #p < 0.05, ##p < 0.01, ###p < 0.001; ####p < 0.0001).

### 3.4. Acarbose prevents metabolic imbalance induced by benzene exposure

To determine whether ACA would mitigate the effects of benzene on metabolic imbalance and hypothalamic inflammation, we fed benzene exposed male mice and their controls with ACA (1,000 mg/kg diet (ppm), as before [39, 42]) for the period of 4 weeks while in the benzene exposure chambers. ACA feeding normalized glucose tolerance, fasted insulin levels and HOMA-IR scores in benzene-treated mice, and they were comparable to control animals (Figure 6A-C). Additionally, ACA treatment reduced serum triglyceride levels, cholesterol, and FFA induced by benzene-exposure to the control levels (Figure 6D-F). 4 weeks of ACA feeding had no effect on body weight or lean and fat body mass in male mice (Supplementary Figure 5). FGF21, a hormone produced by the liver in response to fasting, is elevated by ACA feeding [39]. In support, expression levels of hepatic *Fgf21* were elevated by ACA feeding in control and benzene exposed male mice [39], while expression levels of hepatic *Cyp2e1* were reduced by ACA feeding (Figure 7A, B and Figure 8E). Additionally, ACA feeding inhibited the activation of the hepatic inflammatory genes *Il1b* and *Il6* (Figure 7C and D), and genes associated with gluconeogenesis and lipid metabolism induced by benzene exposure (Figure 7E-K), consistent with the idea of the beneficial effects of ACA on glucose homeostasis [34, 35]. Activation of hypothalamic gliosis was alleviated by the ACA feeding as evidenced by reduced immunoreactivity of GFAP and Iba1 (Figure 8A-D), and normalized expression of inflammatory and ER stress genes in the ARC of ACA-fed benzene exposed male mice (Figure 8F and G). To our knowledge, this is the first demonstration that the anti-diabetes drug Acarbose, can be protective against benzene-induced metabolic imbalance and neuroinflammation, thus providing a broader potential for health promotion.

**Figure 6.**
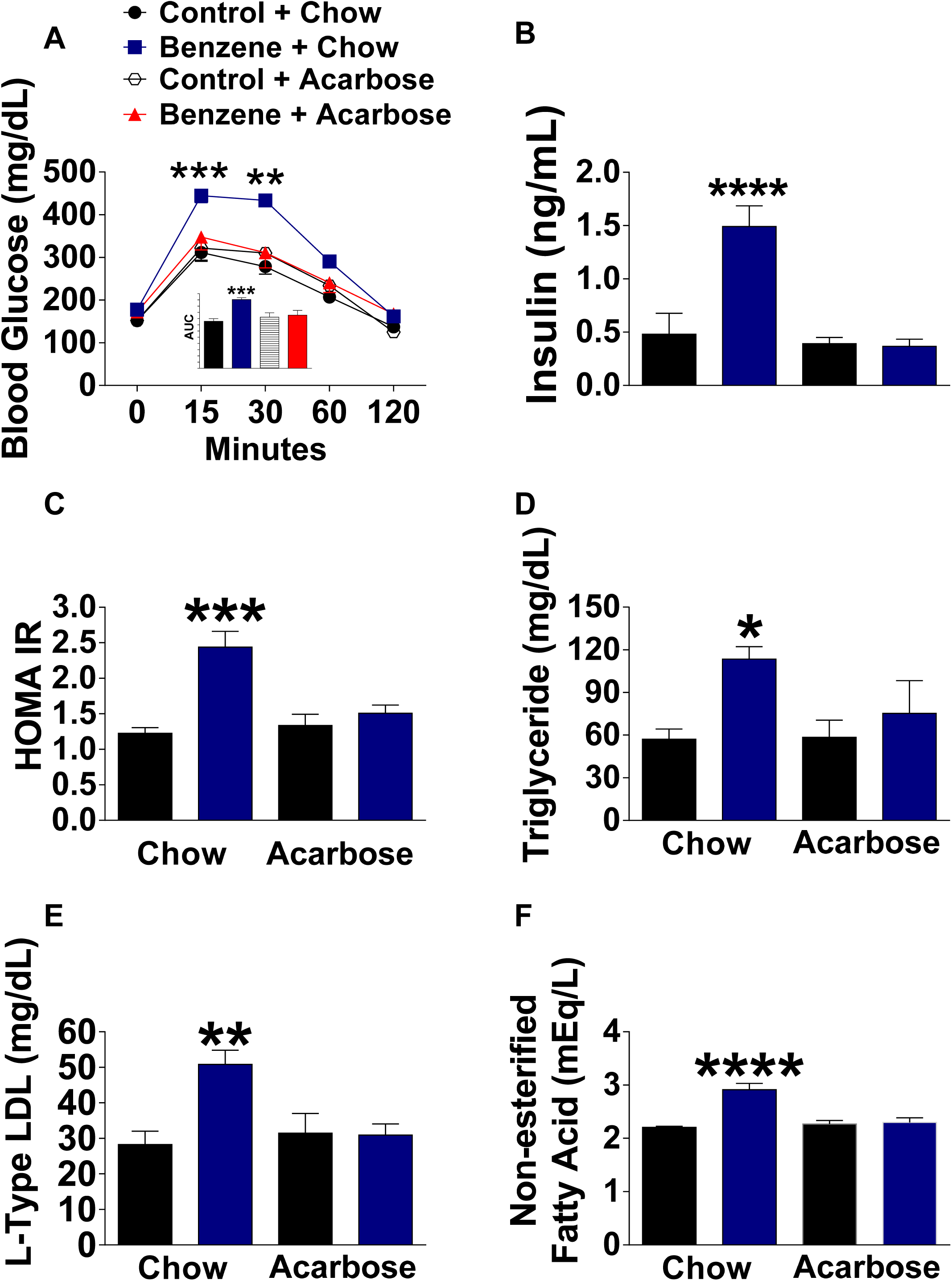
Acarbose prevents metabolic imbalance induced by chronic benzene exposure. Male mice were fed Acarbose (at a concentration of 1,000 mg per kilogram of diet (ppm)) while exposed to benzene (**A**) Glucose tolerance test (effect of treatment: *F*_1.25_ = 10.84, *p*< 0.0029, *η*^2^ = 25.53%; effect of time: *F*_4.100_ = 187.86, *p*< 0.0001, *η*^2^ = 84.32%; diet x time interaction *F*_4.100_ = 3.63, *p* < 0.0082, *η*^2^ = 1.96%; treatment x diet x time interaction *F*_4.100_ = 3.39, *p* < 0.0121, *η*^2^ = 1.52%) and AUC (effect of treatment: *F*_1.37_ = 13.60, *p*< 0.0007, *η*^2^ = 21.51%; treatment x diet interaction *F*_1.37_ = 9.32, *p* < 0.0041, *η*^2^ = 14.74%), (*n* = 6-7); (**B**) Fasted insulin serum levels (effect of treatment: *F*_1.24_ = 64.22, *p*< 0.0001, *η*^2^ = 25.08%; effect of diet: *F*_1.24_ = 96.84, *p* < 0.0001, *η*^2^ = 37.85%; and treatment x diet *F*_1.24_ = 70.87, *p* < 0.0001, *η*^2^ = 27.68%); (**C**) HOMA –IR (effect of treatment: *F*_1.23_ = 15.38, *p*< 0.0006, *η*^2^ = 28.87%; effect of diet: *F*_1.23_ = 5.18, *p* < 0.0324, *η*^2^ = 9.73%; and treatment x diet *F*_1.23_ = 9.72, *p* < 0.0048, *η*^2^ = 18.24%), (*n* = 6-7); (**D**) blood serum levels of triglycerides (effect of treatment: *F*_1.20_ = 6.69, *p*< 0.0176, *η*^2^ = 22.06%); (**E**) LDL (effect of treatment: *F*_1.20_ = 6.90, *p*< 0.0161, *η*^2^ = 17,96% and treatment x diet *F*_1.20_ = 7.59, *p* < 0.0122, *η*^2^ = 19.74%); and (**F**) FFA (effect of treatment: *F*_1.20_ = 17.38, *p*< 0.0004, *η*^2^ = 29.59%; effect of diet: *F*_1.20_ = 11.53, *p* < 0.0028, *η*^2^ = 16.60%; and treatment x diet *F*_1.20_ =17.38, *p* < 0.0001, *η*^2^ = 25.02%), (*n* = 6-7). Data was analyzed with a Two-way ANOVA followed with Newman-Keuls post hoc analysis, (* = *vs* male control; *p < 0.05; **p < 0.01; ***p < 0.001; ****p < 0.0001).

**Figure 7.**
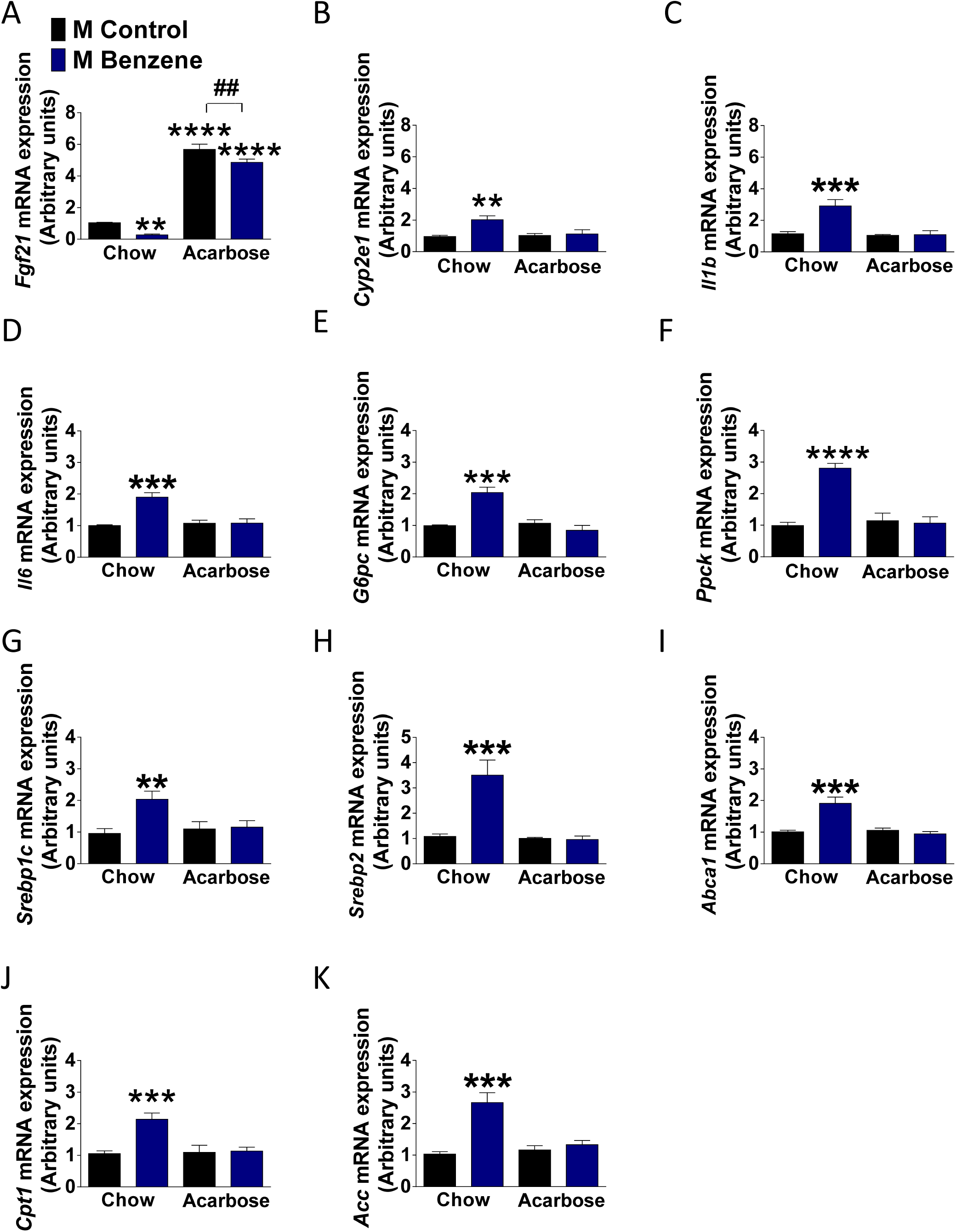
Effect of Acarbose on hepatic gene expression. Hepatic gene expression of (**A**) *Fgf21* (effect of treatment: *F*_1.16_ = 15.16, *p*< 0.0012, *η*^2^ = 2.75%; of diet: *F*_1.16_ = 518.99, *p* < 0.0001, *η*^2^ = 94.33%), (**B**) *Cyp2e1* (effect of treatment: *F*_1.16_ = 8.92, *p*< 0.0087, *η*^2^ = 24.97%; of diet: *F*_1.16_ = 4.58, *p* < 0.0480, *η*^2^ = 12.84%; and treatment x diet *F*_1.16_ =6.20, *p* < 0.0001, *η*^2^ = 17.37%) (**C**) *Il1b* (effect of treatment: *F*_1.20_ = 10.44, *p*< 0.0041, *η*^2^ = 20.29%; of diet: *F*_1.20_ = 11.62, *p* < 0.0028, *η*^2^ = 22.59%; and treatment x diet *F*_1.20_ = 9.38, *p* < 0.0061, *η*^2^ = 18.24%), (**D**) *Il6* (effect of treatment: *F*_1.16_ = 16.44, *p*< 0.0009, *η*^2^ = 27.41%; of diet: *F*_1.16_ = 11.25 *p* < 0.0040, *η*^2^ = 18.76%; and treatment x diet *F*_1.16_ = 16.27, *p* < 0.0009, *η*^2^ = 27.13%), (**E**) *G6pc* (effect of treatment: *F*_1.16_ = 10.39, *p*< 0.0053, *η*^2^ = 14.87%; of diet: *F*_1.16_ = 18.81 *p* < 0.0005, *η*^2^ = 25.90%; and treatment x diet *F*_1.16_ = 24.72, *p* < 0.0001, *η*^2^ =35.35%), (**F**) *Ppck* (effect of treatment: *F*_1.21_ = 25.64, *p*< 0.0001, *η*^2^ = 26.17%; of diet: *F*_1.21_ = 21.20, *p* < 0.0001, *η*^2^ = 21.64 %; and treatment x diet *F*_1.21_ = 30.14, *p* < 0.0001, *η*^2^ = 30.75%), (**G**) *Srebp1c* (effect of treatment: *F*_1.21_ = 6.08, *p*< 0.0223, *η*^2^ = 17.59% and treatment x diet *F*_1.21_ = 4.89, *p* < 0.0381, *η*^2^ = 14.16%), (**H**) *Srepb2* (effect of treatment: *F*_1.21_ = 8.68, *p*< 0.0077, *η*^2^ = 15.52%; of diet: *F*_1.21_ = 10.31, *p* < 0.0038, *η*^2^ = 21.27%; and treatment x diet *F*_1.21_ = 9.33, *p* < 0.0060, *η*^2^ = 18.83%), (**I**) *Abca1* (effect of treatment: *F*_1.21_ = 7.35, *p*< 0.0130, *η*^2^ = 14.51%; of diet: *F*_1.21_ = 9.97, *p* < 0.0047, *η*^2^ = 19.69%; and treatment x diet *F*_1.21_ = 12.34, *p* < 0.0020, *η*^2^ = 24.35%), (**J**) *Cpt1* (effect of treatment: *F*_1.21_ = 10.37 *p*< 0.0041, *η*^2^ = 21.66%; of diet: *F*_1.21_ = 7.56, *p* < 0.0119, *η*^2^ = 15.80%; and treatment x diet *F*_1.21_ = 8.95, *p* < 0.0069, *η*^2^ = 18.68%), and (**K**) *Acc* (effect of treatment: *F*_1.21_ = 15.55 *p*< 0.0007, *η*^2^ = 28.84%; of diet: *F*_1.21_ = 6.99, *p* < 0.0151, *η*^2^ = 12.98%; and treatment x diet *F*_1.21_ = 10.36, *p* < 0.0041, *η*^2^ = 19.22%), (*n* = 7-5 per each group). Data was analyzed with a Two-way ANOVA followed with Newman-Keuls post hoc analysis, (* = *vs* control chow; *p < 0.05; **p < 0.01; ***p < 0.001; ****p < 0.0001) (# = *vs* control acarbose, #p < 0.05, ##p < 0.01, ###p < 0.001; ####p < 0.0001).

**Figure 8.**
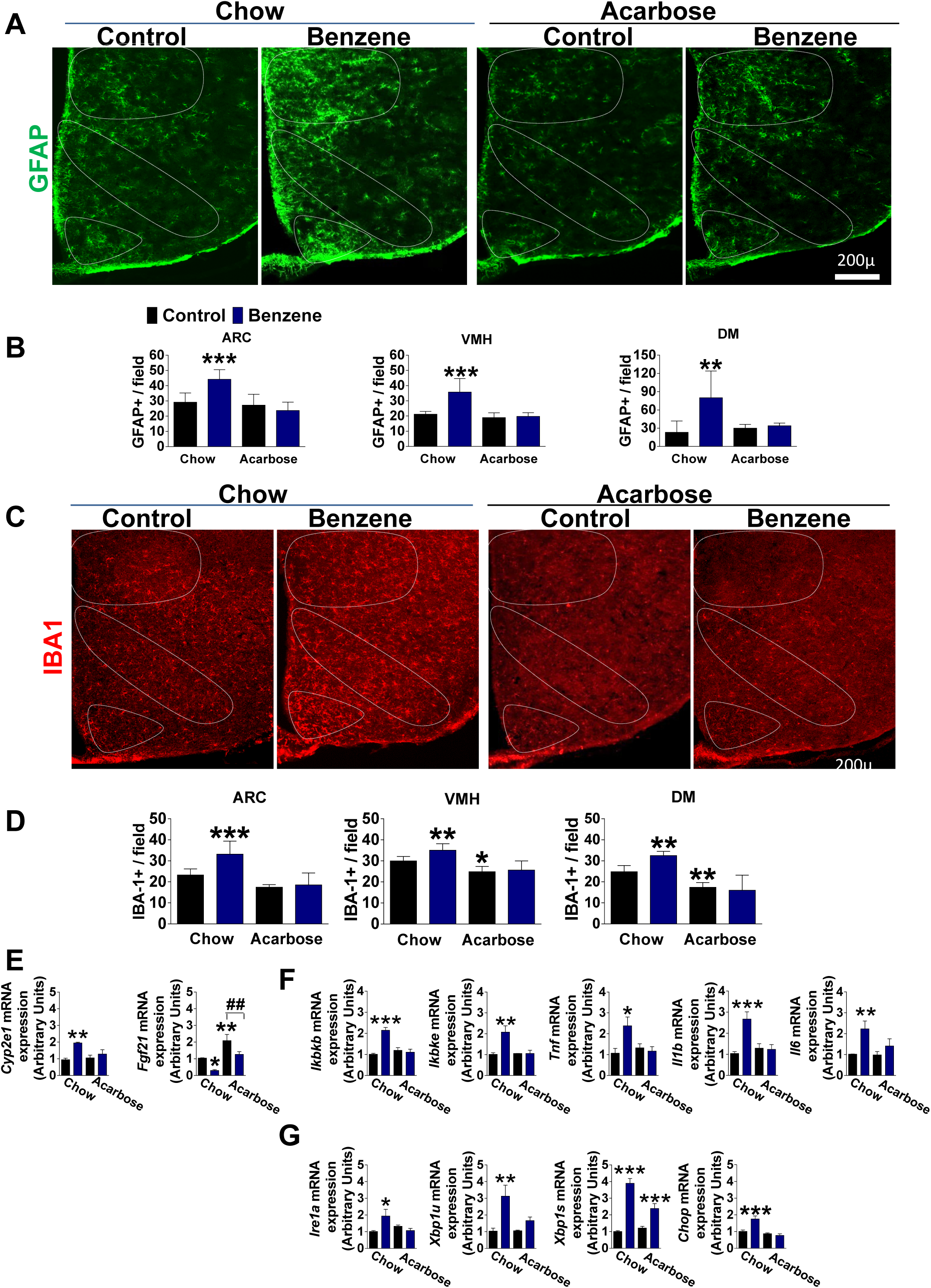
Acarbose prevents hypothalamic gliosis induced by chronic benzene exposure. (**A**) Representative images of astrocytes identified by immunofluorescent detection of GFAP protein in the hypothalamus; (**B**) Quantification of GFAP staining represents number of GFAP positive cells per field (effect of treatment: *F*_1.18_ = 4.48, *p*< 0.0484, *η*^2^ = 8.73%; of diet: *F*_1.18_ = 17.16, *p* < 0.0006, *η*^2^ = 33.44%; and treatment x diet *F*_1.18_ = 11.66, *p* < 0.0031, *η*^2^ = 22.73%), VMH (effect of treatment: *F*_1.18_ = 14.44, *p*< 0.0013, *η*^2^ = 22.48%; of diet: *F*_1.18_ = 20.24, *p* < 0.0002, *η*^2^ = 31.52%; and treatment x diet *F*_1.18_ = 11.54, *p* < 0.0032, *η*^2^ = 17.97%), and DMH (effect of treatment: *F*_1.17_ = 9.73, *p*< 0.0062, *η*^2^ = 25.42%; and treatment x diet *F*_1.17_ = 7.45, *p* < 0.0142, *η*^2^ = 19.47%), (*n* = 7-5). (**C**) Representative images of microglia; (**D**) Quantification of Iba-1 (effect of treatment: *F*_1.18_ = 4.48, *p*< 0.0484, *η*^2^ = 8.73%; of diet: *F*_1.18_ = 17.16, *p* < 0.0006, *η*^2^ = 33.44%; and treatment x diet *F*_1.18_ = 11.66, *p* < 0.0031, *η*^2^ = 22.73%), VMH (effect of treatment: *F*_1.17_ = 4.77, *p*< 0.0432, *η*^2^ =8.96%; of diet: *F*_1.17_ = 28.92, *p* < 0.0001, *η*^2^ = 54.33%), and DMH (effect of diet: *F*_1.18_ = 45.76, *p* < 0.0001, *η*^2^ = 62.25%; and treatment x diet *F*_1.18_ = 6.59, *p* < 0.0193, *η*^2^ = 8.98%) (*n* = 7-5); Gene expression of (**E**) *Cyp2e1* (effect of treatment: *F*_1.12_ = 14.92, *p* < 0.0021, *η*^2^ = 41.70%; and treatment x diet *F*_1.12_ = 5.82, *p* < 0.0328, *η*^2^ = 16.25%) and *Fgf21* (effect of treatment: *F*_1.12_ = 14.53, *p* < 0.0024, *η*^2^ = 28.71%; and of diet *F*_1.12_ = 24.05, *p* < 0.0003, *η*^2^ = 47.51%), (**F**) *Ikbkb* (effect of treatment: *F*_1.12_ = 18.87, *p* < 0.0009, *η*^2^ = 27.38%; and of diet *F*_1.12_ = 12.38, *p* < 0.0042, *η*^2^ = 17.96%; and treatment x diet *F*_1.12_ = 25.69, *p* < 0.0002, *η*^2^ = 37.26%), *Ikbke* (effect of treatment: *F*_1.12_ = 9.14, *p* < 0.0005, *η*^2^ = 24.09%; and of diet *F*_1.12_ = 7.74, *p* < 0.0065, *η*^2^ = 20.39%; and treatment x diet *F*_1.12_ = 0107, *p* < 0.0002, *η*^2^ = 23.92%), *Tnf-α* (effect of treatment: *F*_1.12_ = 6.93, *p* < 0.0218, *η*^2^ = 26.43%), *Il1b* (effect of treatment: *F*_1.12_ = 11.18, *p* < 0.0058, *η*^2^ = 26.43%; and main effect of diet *F*_1.12_ = 12.81, *p* < 0.0037, *η*^2^ = 30.29%), *Il6* (effect of treatment: *F*_1.12_ = 9.45, *p* < 0.0096, *η*^2^ = 36.13%) (*n* =4). (**G**) *Ire1α* (effect of treatment x diet: *F*_1.12_ = 7.45, *p* < 0.0183, *η*^2^ = 31.67%), *Xbp1u* (effect of treatment: *F*_1.12_ = 14.05, *p* < 0.0028, *η*^2^ = 40.84%), *Xbp1s* (effect of treatment: *F*_1.12_ = 16.51, *p* < 0.0001, *η*^2^ = 70.64%; and of diet *F*_1.12_ = 1.69, *p* < 0.0007, *η*^2^ = 2.23%; and treatment x diet *F*_1.12_ = 2.94, *p* < 0.0018, *η*^2^ = 12.60%), *Chop* (effect of treatment: *F*_1.12_ = 7.70, *p* < 0.0016, *η*^2^ = 13.39%; and of diet *F*_1.12_ = 24.11, *p* < 0.0004, *η*^2^ = 41.90%; and treatment x diet *F*_1.12_ = 13.73, *p* < 0.0030, *η*_2_ = 23.87%), (*n* =4). Data was analyzed by Two-way ANOVA followed with Newman-Keuls post hoc analysis, (* = *vs* control chow; *p < 0.05; **p < 0.01; ***p < 0.001; ****p < 0.0001) (# = *vs* control acarbose, #p < 0.05, ##p < 0.01, ###p < 0.001; ####p < 0.0001).

## 4. Discussion

Diabetes is a major public health problem that is approaching epidemic proportions globally, and pollutants in the air or at home have been associated with diabetes [54]. Benzene exposure through the human lifespan is inevitable [6-10]. Drinking contaminated water or eating food that has been exposed to contaminated water, smoking cigarettes, or breathing secondhand cigarette smoke, exposure to e-cigarettes vaping, use of paint or detergent products, vehicle exhaust or gasoline, it’s all part of the daily routine [55]. Our findings demonstrate that chronic benzene exposure induces severe metabolic imbalance associated with central hypothalamic inflammation and ER stress in a sex-specific manner, affecting male but not female mice. Acute benzene exposure robustly activates hypothalamic ER stress via the IRE1α-XBP1 pathway while pharmacological inhibition of the IRE1α alleviates glial inflammatory reaction suggesting that this may be the primary mechanism by which benzene exposure causes changes in metabolic status. Feeding mice anti-diabetes drug Acarbose that also extends mice lifespan, and reduce age-associated hypothalamic inflammation, was sufficient to completely restore central and metabolic imbalance induced by chronic benzene exposure. Collectively, we provide evidence that exposure to benzene, at concentrations relevant to smoking robustly stimulate hypothalamic ER stress and gliosis, which in turn can affect an individual’s susceptibility to metabolic imbalance, and development of T2DM.

Male mice exposed to benzene developed hyperglycemia and hyperinsulinemia followed by elevated expression of hepatic gluconeogenic and lipid metabolism genes. The association between benzene exposure and insulin resistance was reported in a few epidemiological and rodent studies [23, 26, 27, 43, 55, 56]. Studies in humans found that the elimination of benzene is slower in women than in men, likely due to the higher percentage and distribution of body fat tissue [57]. On the other hand, an association between urinary benzene metabolite levels and insulin resistance in elderly adults demonstrated a stronger relationship in men than in women [23]. Furthermore, sex differences were observed upon occupational benzene exposure, particularly effects related to biotransformation of benzene to t,t-MA and hematological parameters [58]. Similarly, benzene exposure rapidly induced hypothalamic and hepatic expression of benzene metabolite, CYP2E1 specifically in male mice, supporting the gender differences in benzene metabolism. Further research is needed to address the physiological reasons underlying the mechanisms by which the sexes differ in their metabolic responses to benzene exposure.

Only male mice show apparent changes in hypothalamic inflammation, ER stress response and peripheral metabolic imbalance due to acute or chronic benzene exposure. While benzene exposure activates hepatic inflammatory genes in both sexes, this effect was not sufficient to induce metabolic imbalance, further supporting the central role of the hypothalamus in benzene induced metabolic imbalance in male mice. Similarly, females are generally more resistant to metabolic imbalance and do not develop hypothalamic inflammation in response to diet-induced obesity, implicating glial responsiveness and activation to this phenomenon [46, 59]. On the other hand, in males, the hypothalamic accumulation of activated microglia is sufficient to trigger metabolic imbalance and susceptibility to obesity [60].

Our data suggest that the hypothalamic IRE1α-XBP1 pathway which is activated during ER stress by benzene exposure can stimulate hypothalamic inflammation since pharmacological inhibition of this pathway by small-molecule inhibitor STF-083010 mitigates glial inflammatory responses in benzene-treated primary glial cells. Impairments in glucose metabolism triggered by benzene exposure were independent of changes in body weight or body composition. In support, short-term hypothalamic ER stress induces systemic insulin resistance independent of body weight or adiposity [61]. Hence, our study suggests a paradigm by which hypothalamic ER stress response induced by benzene exposure can mediate peripheral metabolic imbalance associated with T2D and related problems. These findings complement the recent research which has proposed that exposure to PM_2.5_ induces inflammation via the activation of the IRE1α -XBP1 and ATF6 pathways in zebrafish and mice hepatocytes [62]. Chronic benzene exposure robustly activates hypothalamic ER stress response and, stimulates glial inflammatory reaction highlighting the susceptibility of the hypothalamus to stress and inflammation that lead to changes in metabolic status.

To our knowledge, this is the first direct demonstration that Acarbose, an anti-diabetic drug, can protect against benzene induced metabolic imbalance. Adding Acarbose to chow diet beginning at 4 months of age, was associated with an increase in male (22%) and female (5%) lifespan [38, 39]. This effect was related to increased levels of serum FGF-21, and reduced levels of IGF-1 [39]. We have previously demonstrated that a similar feeding regiment protected mice from age-associated hypothalamic inflammation in a sex-specific manner [42]. While benzene exposure reduced FGF-21 expression, Acarbose increased FGF-21 gene expression in the liver and hypothalamus in male mice. Multiple studies provided evidence that favorable metabolic health impacts of Acarbose are associated with increased production of glucagon-like peptide-1 (GLP-1) by intestinal L cells [63-65]. GLP-1 signaling directly modulates the ER stress response, leading to the promotion of beta-cell survival [66]. Furthermore, GLP-1 production promotes fatty acid oxidation and inhibits fatty acid synthesis [67, 68], which potentially explains our findings showing normalization of serum levels of TG, cholesterol, and FFA upon Acarbose feeding in benzene exposed mice.

Our findings on the protective effects of Acarbose have significant implications for human exposure to benzene. Physiologically, Acarbose imitates the slowing of dietary carbohydrate digestion[69], suggesting that choosing a diet with a low glycemic index or diet in which a slow-digesting carbohydrate predominates, might be a potential strategy for reducing the toxic effects of benzene even at lower levels [70]. Specifically, such a strategy may have the potential for optimizing healthspan for smokers or for people living/working in urban environments with high concentrations of exposure to automobile exhausts.

## Abbreviations

CNS: central nervous system;
ARC: arcuate nucleus of the hypothalamus;
DMH: dorsomedial hypothalamic nucleus;
LHA: lateral hypothalamus;
PVH: paraventricular hypothalamic nucleus;

## 5. Author contributions

LKD, MA, OD, LJBM, PF, IA and AAA, carried out the research and reviewed the manuscript. MS, KZ and UK designed the study and analyzed the data. MS wrote the manuscript and is responsible for the integrity of this work. All authors approved the final version of the manuscript.

## 6. Acknowledgements

This study was supported by American Diabetes Association grant #1-lB-IDF-063, CURES Center Grant (P30 ES020957) and WSU startup funds for MS and UK.

## 8. Data availability

All datasets generated during and/or analyzed during the current study are available from the corresponding author upon request.

## References

1. Kliucininkas, L., et al., Indoor and outdoor concentrations of fine particles, particle-bound PAHs and volatile organic compounds in Kaunas, Lithuania. J Environ Monit, 2011. 13(1): p. 182–91.

2. Briggs, D., Environmental pollution and the global burden of disease. Br Med Bull, 2003. 68: p. 1–24.

3. EPA, Priority pollutants. Washington, D.C., 2012.

4. Tchepel, O., A. Penedo, and M. Gomes, Assessment of population exposure to air pollution by benzene. Int J Hyg Environ Health, 2007. 210(3-4): p. 407–10.

5. Li, L., et al., Pollution characteristics and health risk assessment of benzene homologues in ambient air in the northeastern urban area of Beijing, China. J Environ Sci (China), 2014. 26(1): p. 214–23.

6. Kasemy, Z.A., et al., Environmental and Health Effects of Benzene Exposure among Egyptian Taxi Drivers. J Environ Public Health, 2019. 2019: p. 7078024.

7. Ahola-Olli, A.V., et al., Circulating metabolites and the risk of type 2 diabetes: a prospective study of 11,896 young adults from four Finnish cohorts. Diabetologia, 2019. 62(12): p. 2298–2309.

8. Barros, N., et al., Environmental and biological monitoring of benzene, toluene, ethylbenzene and xylene (BTEX) exposure in residents living near gas stations. J Toxicol Environ Health A, 2019. 82(9): p. 550–563.

9. Wang, Z., et al., The impact of chronic environmental metal and benzene exposure on human urinary metabolome among Chinese children and the elderly population. Ecotoxicol Environ Saf, 2019. 169: p. 232–239.

10. Abdel Maksoud, H.A., et al., Biochemical study on occupational inhalation of benzene vapours in petrol station. Respir Med Case Rep, 2019. 27: p. 100836.

11. Dodson, R.E., et al., Influence of basements, garages, and common hallways on indoor residential volatile organic compound concentrations. Atmospheric Environment, 2008. 42(7): p. 1569–1581.

12. Pankow, J.F., et al., Benzene formation in electronic cigarettes. PLoS One, 2017. 12(3): p. e0173055.

13. Kosmider, L., et al., Cherry-flavoured electronic cigarettes expose users to the inhalation irritant, benzaldehyde. Thorax, 2016. 71(4): p. 376–7.

14. Salviano Dos Santos, V.P., et al., Benzene as a Chemical Hazard in Processed Foods. Int J Food Sci, 2015. 2015: p. 545640.

15. Mansi, A., et al., Low occupational exposure to benzene in a petrochemical plant: modulating effect of genetic polymorphisms and smoking habit on the urinary t,t-MA/SPMA ratio. Toxicol Lett, 2012. 213(1): p. 57–62.

16. Abplanalp, W., et al., Benzene exposure is associated with cardiovascular disease risk. PLoS One, 2017. 12(9): p. e0183602.

17. Duarte-Davidson, R., et al., Benzene in the environment: an assessment of the potential risks to the health of the population. Occup Environ Med, 2001. 58(1): p. 2–13.

18. Wallace, L.A., Major sources of benzene exposure. Environ Health Perspect, 1989. 82: p. 165–9.

19. Bahadar, H., S. Mostafalou, and M. Abdollahi, Current understandings and perspectives on non-cancer health effects of benzene: a global concern. Toxicol Appl Pharmacol, 2014. 276(2): p. 83–94.

20. Brancati, F.L., et al., Diabetes mellitus, race, and socioeconomic status. A population-based study [see comments]. Ann Epidemiol. 1996. Jan. 6: p. 67–73.

21. Fujimoto, W.Y., Overview of non-insulin-dependent diabetes mellitus (NIDDM) in different population groups. Diabet. Med. 1996. Sep. 13: p. S7–10.

22. Shaw, J.E., R.A. Sicree, and P.Z. Zimmet, Global estimates of the prevalence of diabetes for 2010 and 2030. Diabetes Res Clin Pract, 2010. 87(1): p. 4–14.

23. Choi, Y.H., et al., Urinary benzene metabolite and insulin resistance in elderly adults. Sci Total Environ, 2014. 482-483: p. 260–8.

24. Williams, A.D., et al., Ambient Volatile Organic Compounds and Racial/Ethnic Disparities in Gestational Diabetes Mellitus: Are Asian/Pacific Islander Women at Greater Risk? Am J Epidemiol, 2019. 188(2): p. 389–397.

25. Burg, J.R. and G.L. Gist, The National Exposure Registry: analyses of health outcomes from the benzene subregistry. Toxicol Ind Health, 1998. 14(3): p. 367–87.

26. Amin, M.M., et al., Association of benzene exposure with insulin resistance, SOD, and MDA as markers of oxidative stress in children and adolescents. Environ Sci Pollut Res Int, 2018. 25(34): p. 34046–34052.

27. Abplanalp, W.T., et al., Benzene Exposure Induces Insulin Resistance in Mice. Toxicol Sci, 2019. 167(2): p. 426–437.

28. Williams, G., et al., The hypothalamus and the control of energy homeostasis: different circuits, different purposes. Physiol Behav, 2001. 74(4-5): p. 683–701.

29. Timper, K. and J.C. Bruning, Hypothalamic circuits regulating appetite and energy homeostasis: pathways to obesity. Dis Model Mech, 2017. 10(6): p. 679–689.

30. Zhang, X., et al., Hypothalamic IKKbeta/NF-kappaB and ER stress link overnutrition to energy imbalance and obesity. Cell, 2008. 135(1): p. 61–73.

31. Zhang, G., et al., Hypothalamic programming of systemic ageing involving IKK-beta, NF-kappaB and GnRH. Nature, 2013. 497(7448): p. 211–216.

32. Thaler, J.P., et al., Obesity is associated with hypothalamic injury in rodents and humans. J. Clin. Invest, 2012. 122(1): p. 153–162.

33. Liu, C., et al., Central IKKbeta inhibition prevents air pollution mediated peripheral inflammation and exaggeration of type II diabetes. Part Fibre Toxicol, 2014. 11: p. 53.

34. Balfour, J.A. and D. McTavish, Acarbose. An update of its pharmacology and therapeutic use in diabetes mellitus. Drugs, 1993. 46(6): p. 1025–54.

35. DiNicolantonio, J.J., J. Bhutani, and J.H. O’Keefe, Acarbose: safe and effective for lowering postprandial hyperglycaemia and improving cardiovascular outcomes. Open Heart, 2015. 2(1): p. e000327.

36. Sun, W., et al., Comparison of acarbose and metformin therapy in newly diagnosed type 2 diabetic patients with overweight and/or obesity. Curr Med Res Opin, 2016. 32(8): p. 1389–96.

37. Gu, S., et al., Comparison of glucose lowering effect of metformin and acarbose in type 2 diabetes mellitus: a meta-analysis. PLoS One, 2015. 10(5): p. e0126704.

38. Harrison, D.E., et al., Acarbose improves health and lifespan in aging HET3 mice. Aging Cell, 2019. 18(2): p. e12898.

39. Harrison, D.E., et al., Acarbose, 17-alpha-estradiol, and nordihydroguaiaretic acid extend mouse lifespan preferentially in males. Aging Cell, 2014. 13(2): p. 273–82.

40. Garratt, M., et al., Sex differences in lifespan extension with acarbose and 17-alpha estradiol: gonadal hormones underlie male-specific improvements in glucose tolerance and mTORC2 signaling. Aging Cell, 2017. 16(6): p. 1256–1266.

41. Smith, B.J., et al., Changes in the gut microbiome and fermentation products concurrent with enhanced longevity in acarbose-treated mice. BMC Microbiol, 2019. 19(1): p. 130.

42. Sadagurski, M., G. Cady, and R.A. Miller, Anti-aging drugs reduce hypothalamic inflammation in a sex-specific manner. Aging Cell, 2017. 16(4): p. 652–660.

43. De Donno, A., et al., Health Risk Associated with Exposure to PM10 and Benzene in Three Italian Towns. Int J Environ Res Public Health, 2018. 15(8).

44. Lorkiewicz, P., et al., Comparison of Urinary Biomarkers of Exposure in Humans Using Electronic Cigarettes, Combustible Cigarettes, and Smokeless Tobacco. Nicotine Tob Res, 2019. 21(9): p. 1228–1238.

45. Bernauer, U., et al., CYP2E1-dependent benzene toxicity: the role of extrahepatic benzene metabolism. Arch Toxicol, 1999. 73(4-5): p. 189–96.

46. Morselli, E., et al., Hypothalamic PGC-1alpha protects against high-fat diet exposure by regulating ERalpha. Cell Rep, 2014. 9(2): p. 633–45.

47. Morselli, E., et al., A sexually dimorphic hypothalamic response to chronic high-fat diet consumption. Int J Obes (Lond), 2016. 40(2): p. 206–9.

48. Kapoor, N., et al., Differences in sensitivity of cultured rat brain neuronal and glial cytochrome P450 2E1 to ethanol. Life Sci, 2006. 79(16): p. 1514–22.

49. Tokutake, Y., et al., IRE1-XBP1 Pathway of the Unfolded Protein Response Is Required during Early Differentiation of C2C12 Myoblasts. Int J Mol Sci, 2019. 21(1).

50. Zhang, K. and R.J. Kaufman, From endoplasmic-reticulum stress to the inflammatory response. Nature, 2008. 454(7203): p. 455–62.

51. Qiu, Q., et al., Toll-like receptor-mediated IRE1alpha activation as a therapeutic target for inflammatory arthritis. EMBO J, 2013. 32(18): p. 2477–90.

52. He, Y., et al., Emerging roles for XBP1, a sUPeR transcription factor. Gene Expr, 2010. 15(1): p. 13–25.

53. Kim, S., et al., Endoplasmic reticulum stress-induced IRE1alpha activation mediates cross-talk of GSK-3beta and XBP-1 to regulate inflammatory cytokine production. J Immunol, 2015. 194(9): p. 4498–506.

54. Dong, G., et al., Effect of Social Factors and the Natural Environment on the Etiology and Pathogenesis of Diabetes Mellitus. Int J Endocrinol, 2019. 2019: p. 8749291.

55. Canney, S., EPA regulatory update. Occup Health Saf, 2002. 71(11): p. 58–60, 84.

56. Thiering, E., et al., Long-term exposure to traffic-related air pollution and insulin resistance in children: results from the GINIplus and LISAplus birth cohorts. Diabetologia, 2013. 56(8): p. 1696–704.

57. Sato, A., et al., Kinetic studies on sex difference in susceptibility to chronic benzene intoxication--with special reference to body fat content. Br J Ind Med, 1975. 32(4): p. 321–8.

58. Moro, A.M., et al., Biomonitoring of gasoline station attendants exposed to benzene: Effect of gender. Mutat Res, 2017. 813: p. 1–9.

59. Dorfman, M.D., et al., Sex differences in microglial CX3CR1 signalling determine obesity susceptibility in mice. Nat Commun, 2017. 8: p. 14556.

60. Valdearcos, M., et al., Microglia dictate the impact of saturated fat consumption on hypothalamic inflammation and neuronal function. Cell Rep, 2014. 9(6): p. 2124–38.

61. Purkayastha, S., et al., Neural dysregulation of peripheral insulin action and blood pressure by brain endoplasmic reticulum stress. Proc Natl Acad Sci U S A, 2011. 108(7): p. 2939–44.

62. Zhang, Y., et al., Developmental toxicity induced by PM2.5 through endoplasmic reticulum stress and autophagy pathway in zebrafish embryos. Chemosphere, 2018. 197: p. 611–621.

63. Qualmann, C., et al., Glucagon-like peptide 1 (7-36 amide) secretion in response to luminal sucrose from the upper and lower gut. A study using alpha-glucosidase inhibition (acarbose). Scand J Gastroenterol, 1995. 30(9): p. 892–6.

64. DeLeon, M.J., et al., Glucagon-like peptide-1 response to acarbose in elderly type 2 diabetic subjects. Diabetes Res Clin Pract, 2002. 56(2): p. 101–6.

65. Wang, N., et al., Associations between changes in glucagon-like peptide-1 and bodyweight reduction in patients receiving acarbose or metformin treatment. J Diabetes, 2017. 9(8): p. 728–737.

66. Yusta, B., et al., GLP-1 receptor activation improves beta cell function and survival following induction of endoplasmic reticulum stress. Cell Metab, 2006. 4(5): p. 391–406.

67. Ben-Shlomo, S., et al., Glucagon-like peptide-1 reduces hepatic lipogenesis via activation of AMP-activated protein kinase. J Hepatol, 2011. 54(6): p. 1214–23.

68. Galsgaard, K.D., et al., Glucagon Receptor Signaling and Lipid Metabolism. Front Physiol, 2019. 10: p. 413.

69. Pham, H., et al., A randomized, crossover study of the acute effects of acarbose and gastric distension, alone and combined, on postprandial blood pressure in healthy older adults. BMC Geriatr, 2019. 19(1): p. 241.

70. Jenkins, D.J., Lente carbohydrate: a newer approach to the dietary management of diabetes. Diabetes Care, 1982. 5(6): p. 634–41.

71. Sadagurski, M., et al., Human IL6 enhances leptin action in mice. Diabetologia, 2010. 53(3): p. 525–535.

72. Sadagurski, M., et al., Growth hormone modulates hypothalamic inflammation in long-lived pituitary dwarf mice. Aging Cell, 2015. 14(6): p. 1045–54.

73. Pool, M., et al., NeuriteTracer: a novel ImageJ plugin for automated quantification of neurite outgrowth. J Neurosci Methods, 2008. 168(1): p. 134–9.

74. Longair, M.H., D.A. Baker, and J.D. Armstrong, Simple Neurite Tracer: open source software for reconstruction, visualization and analysis of neuronal processes. Bioinformatics, 2011. 27(17): p. 2453–4.

75. Papandreou, I., et al., Identification of an Ire1alpha endonuclease specific inhibitor with cytotoxic activity against human multiple myeloma. Blood, 2011. 117(4): p. 1311–4.

76. Chien, W., et al., Selective inhibition of unfolded protein response induces apoptosis in pancreatic cancer cells. Oncotarget, 2014. 5(13): p. 4881–94.

