## Supplemental material for "Acarbose Protects From Central and Peripheral Metabolic Imbalance Induced by Benzene Exposure"

### Supplementary Methods

#### Flow Cytometry Analysis

For identification of the leukocyte population, single-cell suspensions were stained with fluorescent-labeled antibodies to detect myeloid cells ($CD45^+CD11b^+$), neutrophils ($CD45^+CD11b^+Ly6G^+$), macrophages ($CD45^+$ $CD11b^+$ $Ly6G^-$ $Ly6G^{low}$), monocytes ($CD45^+$ $CD11b^+$ $Ly6G^-$ $Ly6G^{high}$), and lymphocytes (SSC-$A^-$ $CD45^{high}$). Leukocyte cell populations were incubated with 0.5 µL Fc-blocking agent (anti-mouse CD 16/32, Biolegend, San Diego, CA) for 10 minutes at RT prior to the addition of the multi-color antibody panel. Fluorescent minus one (FMO) controls were used to differentiate between positively and negatively stained populations. Compensation was performed using BD Comp Beads (BD Biosciences, San Jose, CA) to create single-color controls of each antibody. All antibodies were obtained from Biolegend (San Diego, CA) unless otherwise stated. FACS analyses were performed on a BD LSR II utilizing the services of the microscopy, imaging, and cytometry core laboratory (MICR), at Wayne State University, Detroit, MI, and data were analyzed with FlowJo software (FlowJo, LLC).

#### Urinary Metabolite analysis

At the end of the benzene exposure, urine was collected and centrifuged at 400xg for 5 min and the supernatant stored at −80°C. 20 µL of urine was diluted into 70 µL of water containing 10% formic acid followed and 10 µL of a solution containing 1 ppm of $^{13}C_6$-t,t-MA. 20 µL of the sample was injected and chromatographed using a Shimadzu Nexera X2 UPLC system with a 100 mm x 2.05 mm x 1.7 um Waters BEH C18 Column fitted with a 0.2 µm filter in a 50 °C oven. Mobile phase A was 0.1% acetic acid in water and mobile phase B was Methanol. The target analyte is highly water-soluble and chromatography was achieved by a slight ramp from 0% B to 10% B over 2 minutes followed by a ramp to 25% B at 4 minutes. This was followed by a rinse and equilibration cycle for a total run time of 10 minutes. The retention time of t,t-

MA was found to be 2.6 minutes. A Shimadzu 8040 LCMSMS was used in MRM mode to analyze for the transition 141 to 97 for t,t-MA and 147 to 102 for $^{13}C_6$- t,t-MA in negative ionization mode. The detection limit was found to be approximately 0.05 ppm with a signal to noise ratio of about 2. Seven individually prepared replicates gave an average calculated concentration of 0.043 ppm with a standard deviation of 0.004 ppm. LOQ is defined as 10 standard deviations from the mean and was found to be 0.09 ppm.Calibration was done by spiking synthetic urine diluent (SigMatrix Urine) at the calibration levels 0.05, 0.1, 0.5, 1.0, 5.0, 10.0, 50.0, 100.0. Calibrants were prepared identically to the samples and extremely high accuracy was found fitting a quadratic, internal standard normalized, 1/C weighted calibration curve. The result was normalized by the control samples.

**SUPPLEMENTARY FIGURE LEGENDS:**

**Supplementary Figure 1.** (**A**) Image of FlexStream system used for the benzene exposure; Effect of benzene exposure on (**B**) fat mass (g) and (**C**) lean mass (g) measured by magnetic resonance imaging (echoMRI), (**D**) blood cell number of neutrophils (Neu), macrophage/monocyte (Macro/Mono), lymphocyte (Lympho) and total cells. Urine levels of (**E**) benzene metabolite levels, t,t-MA ($t$ =5.569, $p < 0.0014$, $n = 4$), and (**F**) *Cyp2e1* expression in the liver ($t$ =3.282, $p < 0.0168$, $n = 4$). Data are expressed as the mean ± SEM. (* p < 0.05 vs control).

**Supplementary Figure 2.** Effect of chronic benzene exposure on insulin tolerance test (ITT) of (**A**) males (M) and (**B**) female (F). Data are expressed as the mean ± SEM.

**Supplementary Figure 3.** Effects of HFD feeding on (**A**) blood cell number of neutrophils (Neu), macrophage/monocyte (Macro/Mono), lymphocyte (Lympho) and total cells; bodyweight of (**B**) male (M) and (**C**) female (F); echoMRI of (**D**) fat mass (g) and (**E**) lean mass (g) of male, (**F**) fat mass (g) and (**G**) lean mass (g) of female. Data are expressed as the mean ± SEM. * p < 0.05 vs control.

**Supplementary Figure 4.** Effect of acute benzene exposure of inflammatory genes expression (**A**) *Ikbkb* ($t$ =5.505, $p < 0.0015$, $n = 4$), (**B**) *Ikbke* ($t$ =2.930, $p < 0.0263$, $n = 4$), (**C**) *Tnf* ($t$ =3.788, $p < 0.0091$, $n = 4$), (**D**) *Il1b* ($t$ =2.813, $p < 0.0306$, $n = 4$), (**E**) *Il6* ($t$ =19.55, $p < 0.0001$, $n = 4$), (**F**) *Cyp2e1* in the ARC. Data are expressed as the mean ± SEM. * p < 0.05 vs control.

**Supplementary Figure 5.** Effects of Acarbose on (**A**) body weight; (**B**) echoMRI of fat mass (g); and (**C**) lean mass (g). Data are expressed as the mean ± SEM. * p < 0.05 vs control.

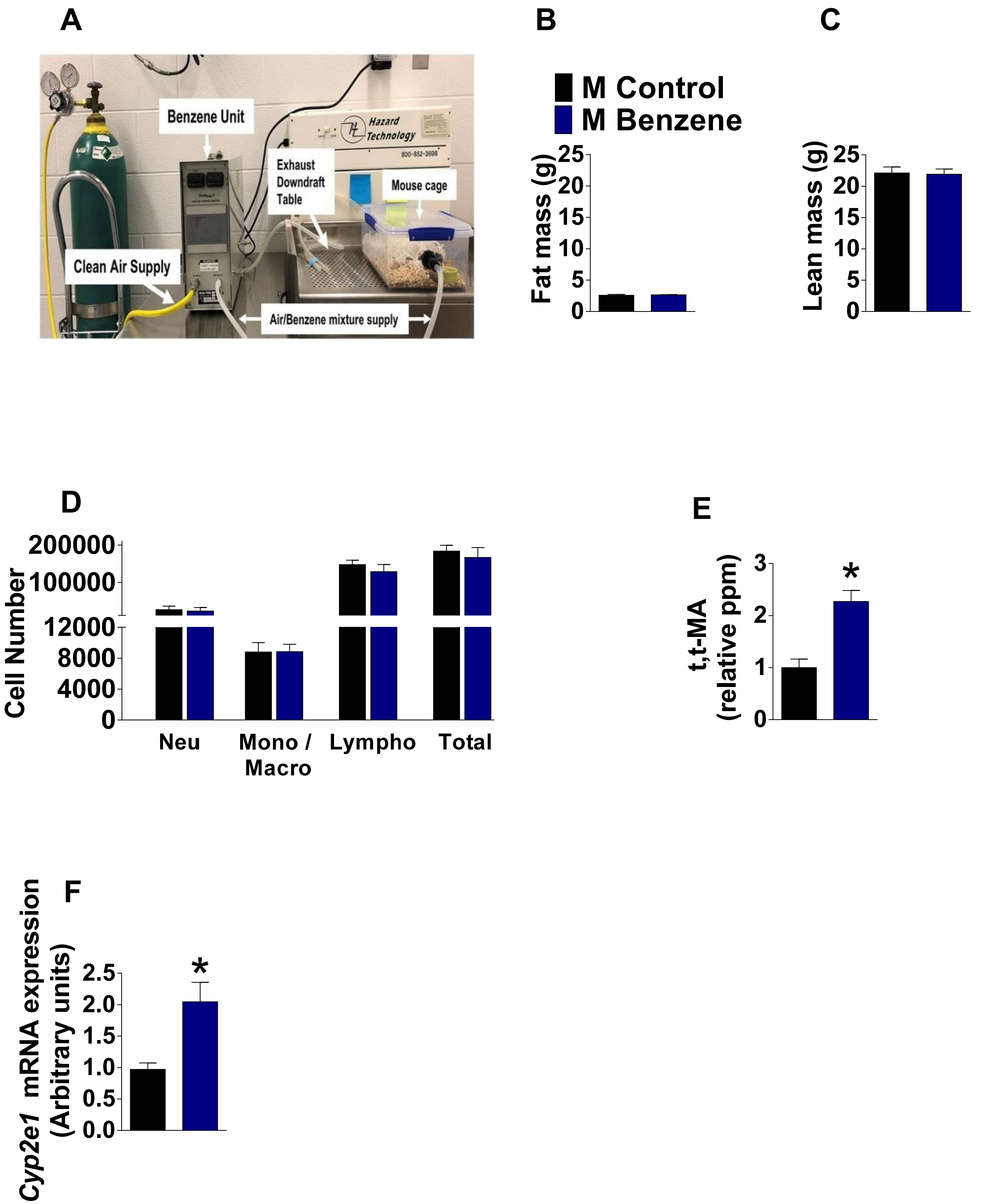


Supplementary Figure 1

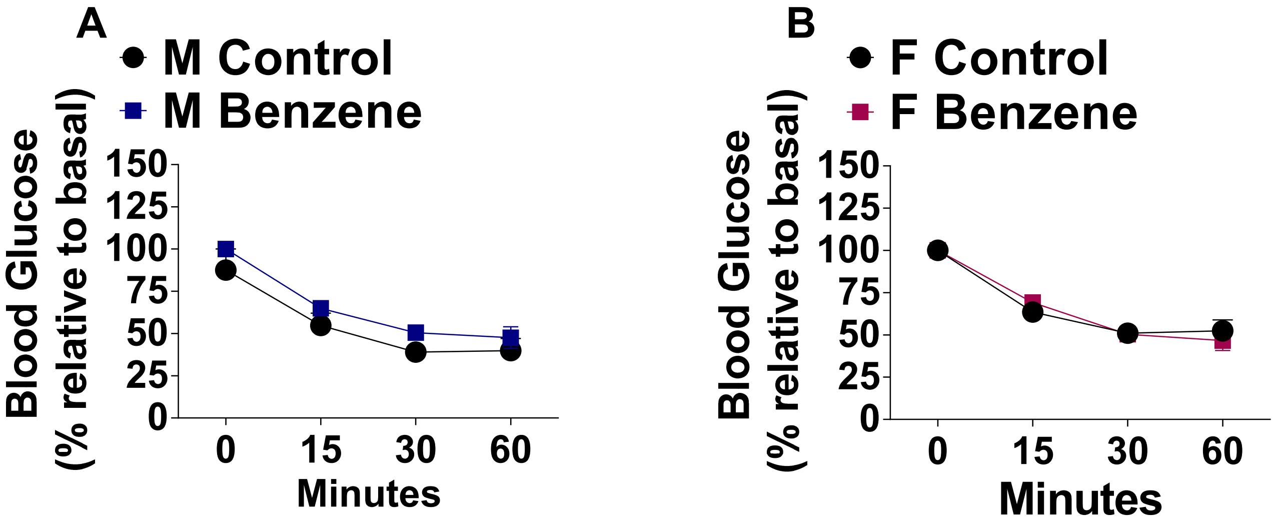


Supplementary Figure 2

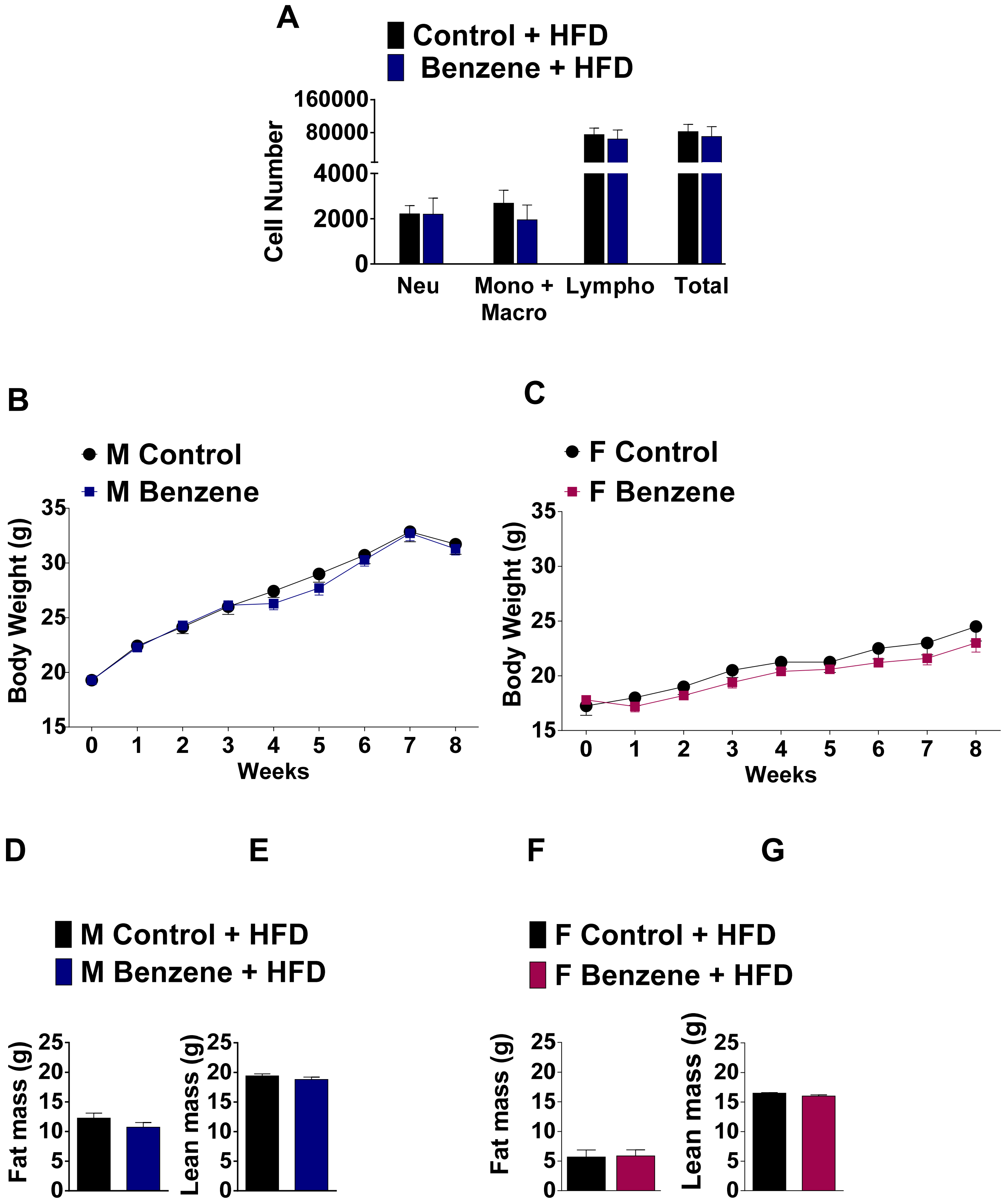


Supplementary Figure 3

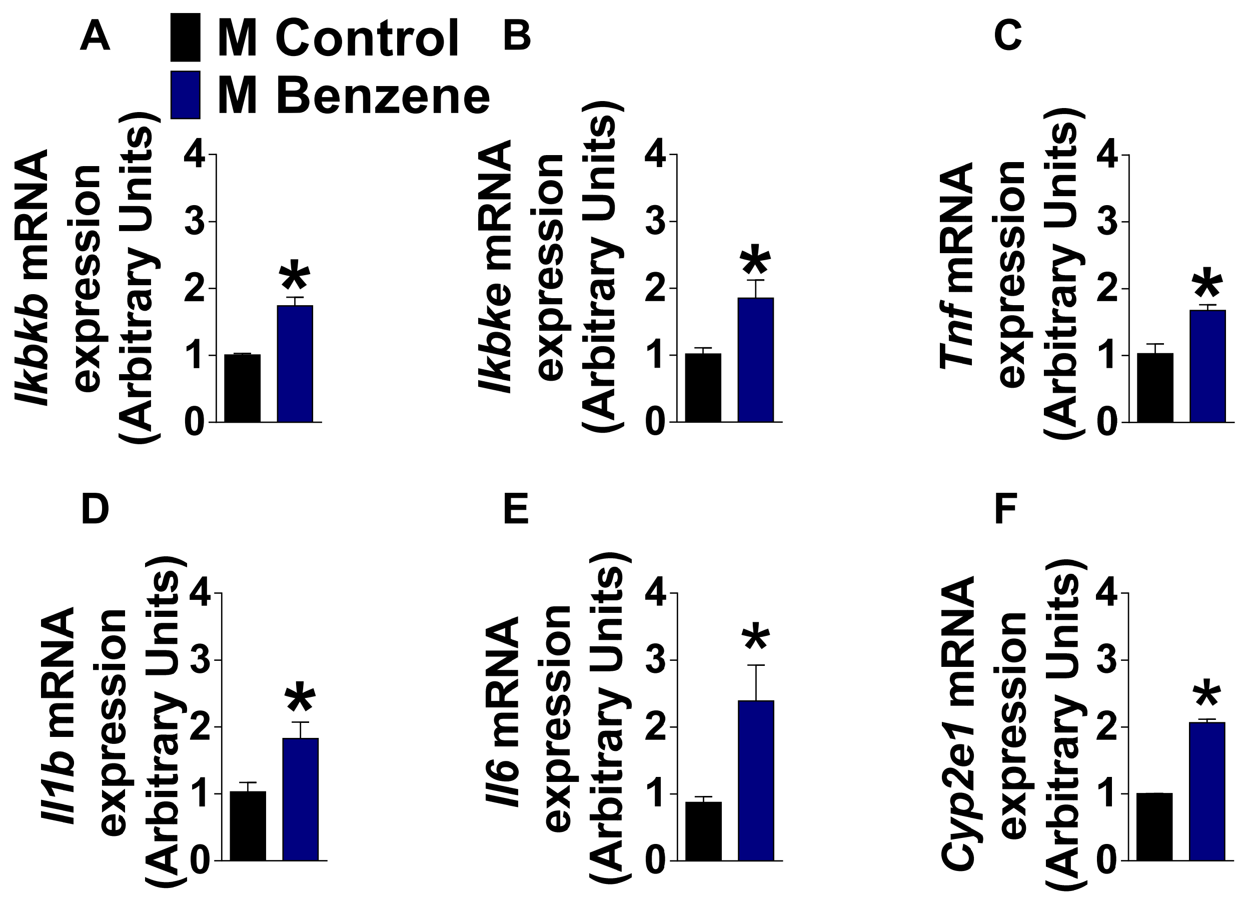


Supplementary Figure 4

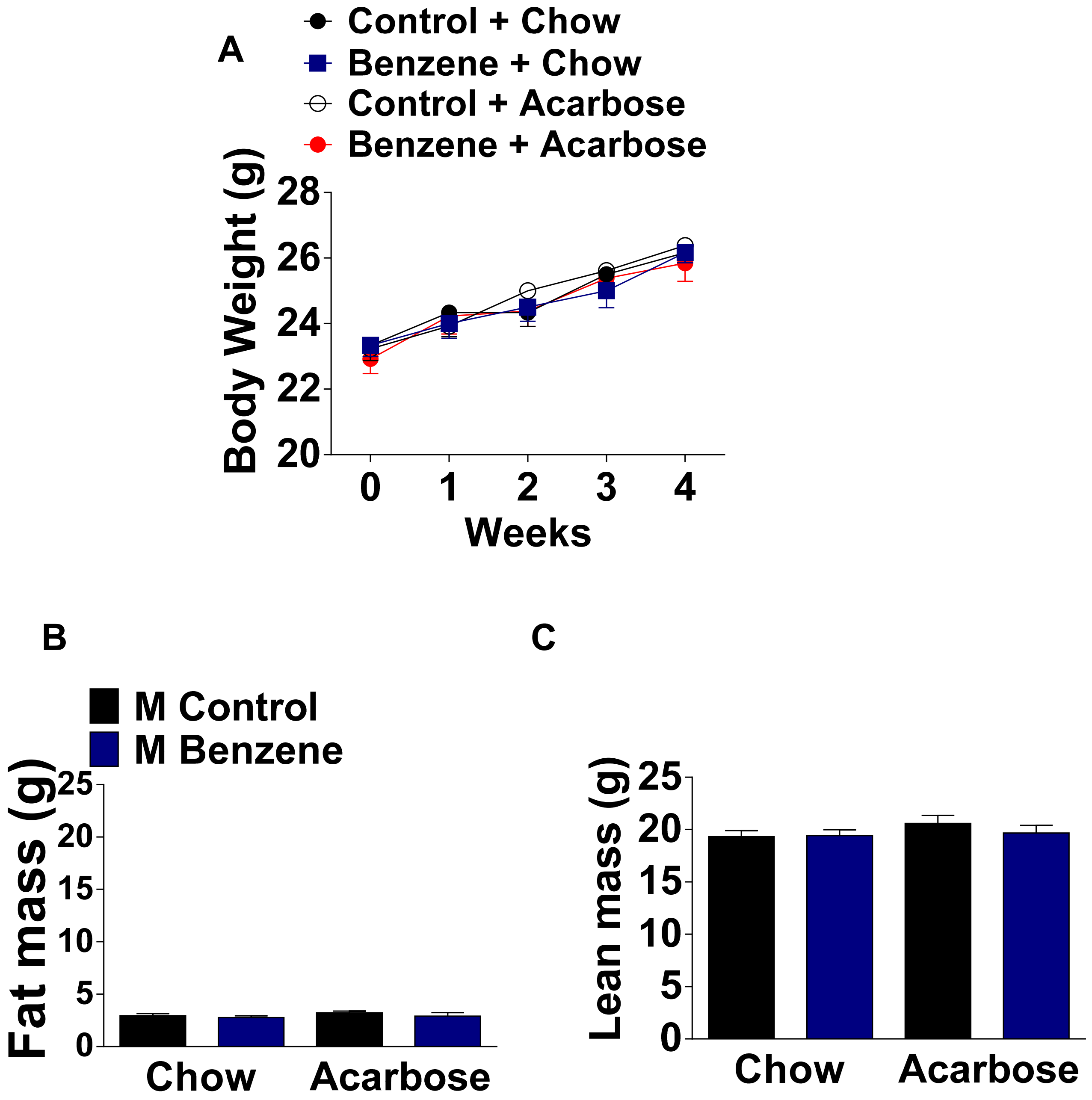


Supplementary Figure 5
